# Assessing the combined effects of forest management and climate change on carbon and water fluxes in European beech forests

**DOI:** 10.1101/2024.08.20.608827

**Authors:** Vincenzo Saponaro, Miquel De Cáceres, Daniela Dalmonech, Ettore D’Andrea, Elia Vangi, Alessio Collalti

## Abstract

The consequences of climate change continue to threaten European forests, particularly for species located at the edges of their latitudinal and altitudinal ranges. While extensively studied in Central Europe, European beech forests require further investigation to understand how climate change will affect these ecosystems in Mediterranean areas. Proposed silvicultural options increasingly aim at sustainable management to reduce biotic and abiotic stresses and enhance these forest ecosystems’ resistance and resilience mechanisms. Process-based models (PBMs) can help us to simulate such phenomena and capture early stress signals while considering the effect of different management approaches. In this study, we focus on estimating sensitivity of two state-of-the-art PBMs forest models by simulating carbon and water fluxes at the stand level to assess productivity changes and feedback resulting from different climatic forcings as well as different management regimes. We applied the 3D-CMCC-FEM and MEDFATE forest models for carbon (C) and water (H_2_O) fluxes in in two sites of the Italian peninsula, Cansiglio in the north and Mongiana in the south, under managed vs. unmanaged scenarios and under current climate and different climatic scenarios (RCP4.5 and RCP8.5). To ensure confidence in the models’ results, we preliminary evaluated their performance in simulating C and H_2_O flux in three additional beech forests of the FLUXNET network along a latitudinal gradient spanning from Denmark to central Italy. The 3D-CMCC-FEM model achieved R² values of 0.83 and 0.86 with RMSEs of 2.53 and 2.05 for C and H_2_O fluxes, respectively. MEDFATE showed R² values of 0.76 and 0.69 with RMSEs of 2.54 and 3.01. At the Cansiglio site in northern Italy, both models simulated a general increase in C and H_2_O fluxes under the RCP8.5 climate scenario compared to the current climate. Still, no benefit in managed plots compared to unmanaged ones, as the site does not have water availability limitations, and thus, competition for water is low. At the Mongiana site in southern Italy, both models predict a decrease in C and H_2_O fluxes and sensitivity to the different climatic forcing compared to the current climate; and an increase in C and H_2_O fluxes when considering specific management regimes compared to unmanaged scenarios. Conversely, under unmanaged scenarios plots are simulated to experience first signals of mortality prematurely due to water stress (MEDFATE) and carbon starvation (3D-CMCC-FEM) scenarios. In conclusion, while management interventions may be considered a viable solution for the conservation of beech forests under future climate conditions at moister sites like Cansiglio, in drier sites like Mongiana conservation may not lie in management interventions alone.

## 1. Introduction

Predicting the future evolution of European forests is essential to continue to benefit from the ecosystem services they provide for human well-being. Forests offer, for instance, climate change mitigation through their ability to store atmospheric carbon dioxide in biomass and soil (Augusto and Bo, 2022; Pan et al., 2024). In 2020, the European Green Deal prioritized the vital role of forests and the forestry sector in attaining sustainability objectives, such as promoting sustainable forest management, enhancing forest resilience, and climate change mitigation (European Commission, 2021). Technological advances and studies of forest ecosystem responses to management practices continue to promote the evolution of strategies that maintain or enhance forest ecosystem services, such as promoting biological diversity, water resources, soil protection, or carbon sequestration (Pukkala, 2016). Different forest management systems have been adopted in Europe over the years (e.g., clear-cutting or shelterwood) depending, among others, on the wood product desired, the stand age, and structure (Brunet et al., 2010).

Forest management can be a key element in mitigating the effects of climate warming, maintaining the current primary productivity and the current distribution of tree species, or altering forest composition to promote more suited and productive species (Nolè et al., 2015; Bosela et al., 2016). Indeed, the carbon sequestration capacity and productivity of forests are dependent, primarily, on species composition, site conditions as well as on stand age (Rötzer et al., 2010; Vangi et al., 2024a, b), which are affected by past and present forest management activities. According to Collalti et al. (2018) and Dalmonech et al. (2022), monospecific forests in Europe would appear unable to further increase current rates of carbon storage and biomass production under future climate scenarios, considering current management practices, but at the same time demonstrating that managing under Business as Usual (BAU) practices still allows forests to accumulate biomass at higher rates compared to stands left to develop undisturbed.

European beech (*Fagus sylvatica* L.) is an important deciduous tree species widely distributed in Europe, from southern Scandinavia to Sicily and Spain to northwest Turkey (Durrant et al., 2016). In Italy, according to the National Forest Inventory (INFC, 2015), beech forests cover a total area of 1,053,183 hectares, accounting for about 11.7% of the country’s overall forested land. European beech forests demonstrate susceptibility to temperature and precipitation fluctuations. For instance, a warmer environment and less precipitation are forcing shifts in distribution area or the onset of loss of canopy greenness (Axer et al., 2021; Noce et al., 2017, 2023; Zuccarini et al., 2023; Rezaie et al., 2018). According to Skrk et al. (2023), the decline in growth of the beech forests primarily occurs in the dry and warm marginal conditions prevalent near the geographical edge of its distribution with a sub-Mediterranean climatic regime, posing a threat to the survival of beech populations in those areas. However, tree ring analyses have also revealed an unexpected increase in growth in the south Mediterranean region of Albania and Macedonia beech forests at the end of the 20^th^ century, challenging the presumed decline of forest ecosystems due to drought (Tegel et al., 2014). Puchi et al. (2024) additionally shed light on the susceptibility to extreme drought events of beech forests found at higher latitudes compared to those found at lower latitudes in the Italian peninsula by highlighting an increase, for the latter, in growth related to the abundance of precipitation. In this context, it is important to minimize the uncertainty surrounding the response of the carbon, water, and energy cycles within beech forest ecosystems, especially as they have been shown to adapt to varying environmental drivers (Deb Burman et al., 2024).

Process-based models (PBMs) are useful tools for studying forest dynamics, as well as water (H_2_O) and carbon (C) use efficiency, and carbon stocks as key variables of forest mitigation potential (Vacchiano et al., 2012; Pilli et al., 2022; Testolin et al., 2023; Morichetti et al., 2024). Forest modelling has been widely used by forest ecologists for tackling numerous applied research questions, and the field is continuously evolving to improve process representation to achieve higher realism and predictive capacity under warmer climate and forest management scenarios (Riviere et al., 2020; Kimmins et al., 2008; Nolè et al., 2013; Maréchaux et al., 2021). By comparing the predictive performance of different models under current environmental conditions, it is possible to gain confidence in their predictions of future trends and make informed decisions in forest ecosystem management and planning processes (Huber et al., 2013; Mahnken et al., 2022).

The main goal of the present study is to evaluate the impact of forest management regimes and climate change scenarios on European beech forests using two state-of-the-science PBMs: 3D-CMCC-FEM (Collalti et al., 2014) and MEDFATE (De Cáceres et al., 2023). More specifically, the study aims to provide deeper insights into the carbon (C) and water (H_2_O) fluxes of this species under varying management practices and changing environmental conditions. The MEDFATE model is capable of simulating the complex water dynamics linking the soil-vegetation-atmosphere continuum. The performance of MEDFATE in simulating soil moisture dynamics and plant transpiration has been extensively evaluated across various scales and different stand structures, particularly in Mediterranean environments (De Cáceres et al., 2015, 2021; Sánchez-Dávila et al., 2024). Complementarily, the 3D-CMCC-FEM model has been extensively validated and shown to effectively capture the spatial and temporal variability of carbon and water fluxes, while accounting for ecological heterogeneity and integrating forest management practices across a wide range of scales (Collalti et al., 2018; Dalmonech et al., 2022, 2024; Mahnken et al., 2022).

Since the study sites vary in terms of environmental factors that can affect gross primary productivity (GPP), as well as latent heat (LE), which are the two variables considered in this analysis, the use of two PBMs can provide the highest reliability in capturing the complex dynamics of these variables under diverse environmental conditions. By leveraging the strengths of both models, we can achieve a more robust and comprehensive understanding of C and H_2_O across the different sites. Specifically, we tested: (i) to what extent different forest management options can influence C and H_2_O fluxes under the present-day climate; and, (ii) how harsher climate conditions may affect the C and H_2_O fluxes under different management options. To gain confidence on the models predictive capacity we preliminary parameterized and evaluated models’ performances for C and H_2_O fluxes at three forest stands dominated by beech forests: the Sorø (DK-Sor), Hesse (FR-Hes), and Collelongo (IT-Col) sites, which are included in the PROFOUND Database (PROFOUND DB) (Reyer et al., 2020a, b) and makes part of the FLUXNET Network (Pastorello et al., 2020). To address the questions, we assessed the C and H_2_O fluxes at two target and independent beech forest sites in Italy (Cansiglio and Mongiana) by simulating their development under various management options and evaluating their (model) sensitivity to current and more severe climate conditions.

## 2. Material and Methods

### 2.1 3D-CMCC-FEM model

The 3D-CMCC-FEM v.5.6 (‘*Three-Dimensional – Coupled Model Carbon Cycle – Forest Ecosystem Module*’) (Collalti et al., 2024, and references therein; Marconi et al., 2017; Dalmonech et al., 2022, 2024; Vangi et al., 2024a,b; Morichetti et al., 2024) is a C-based, eco-physiological, biogeochemical and biophysical model. The model simulates C and H_2_O fluxes occurring within forest ecosystems daily, monthly, or annually, depending on the processes to simulate, with a common spatial scale of one hectare (Collalti et al., 2016). Photosynthesis is simulated using the biochemical model of Farquhar–von Caemmerer–Berry (Farquhar et al., 1980), integrating the sunlit and shaded leaves of the canopy (De Pury and Farquhar, 1997). For the temperature dependence of the Michaelis-Menten coefficient for Rubisco and the CO_2_ compensation point without mitochondrial respiration, the model adopts the parameterization described in Bernacchi et al. (2001, 2003). The net balance at the autotrophic level is represented by net primary production in eq 1:

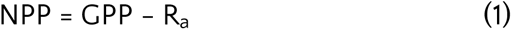

where R_a_ includes both maintenance respiration (R_m_) and growth respiration (R_g_). When R_m_ exceeds GPP, resulting in a negative NPP, the trees utilize their non-structural carbon reserves (NSC) (i.e., soluble sugars and starch, undistinguished) to meet the carbon demand (Merganičová et al., 2019; Collalti et al., 2020a). In deciduous trees, NSC is used to create new leaves during the bud-burst phase, replenishing during the growing season under favourable photosynthetic conditions, and finally remobilising to other tissues to prepare trees for dormancy at the end of the growth phase. The model assumes that NSC reserves are actively mobilized to meet metabolic demands during periods of stress or carbon deficits, such as drought. For instance, during periods of negative carbon balance, the model allocates stored NSC to sustain key physiological processes (e.g., maintenance respiration, leaf and fine root formation). The allocation scheme ensures that NSC replenishment is prioritized before supporting growth demands (i.e., wood growth), consistent with evidence showing that carbon flows are first directed toward restoring NSC reserves until critical thresholds are reached (Hartmann & Trumbore, 2016). Replenishment of non-structural carbon reserves is essential to achieve the minimum safety threshold (i.e., 11% of sapwood dry mass for deciduous trees; Schwalm and Ek, 2004). Failure to meet these thresholds may trigger at first remobilization from leaves and fine root and subsequently to defoliation mechanisms, while complete depletion of reserves (e.g., during prolonged stress periods) could lead to the death of the entire cohort of trees through carbon starvation. In 3D-CMCC-FEM stomatal conductance g_s_ is calculated using the Jarvis equation (Jarvis, 1976). The equation includes a species-specific parameter g_s max_ (i.e., maximum stomatal conductance) controlled by factors such as light, atmospheric CO_2_ concentration, air temperature, soil water content, vapour pressure deficit (VPD), and stand age (Collalti et al., 2019). According to Waring and Running (2007) and Monteith and Unsworth (2008), the Penman-Monteith equation is used to calculate the latent heat (LE) fluxes of evaporation as a function of incoming radiation, VPD, and conductances at a daily scale, summing up the canopy, soil, and snow (if any) latent heat flux expressed as W m^-2^ or MJ m^-2^ time^-1^

The 3D-CMCC-FEM accounts for forest stand dynamics, including growth, competition for light, and tree mortality under different climatic conditions, considering both the CO_2_ fertilization effects and temperature acclimation (Collalti et al., 2018, 2019; Kattge and Knorr, 2007). Several mortality routines are considered in the model, such as age-dependent mortality, background mortality (stochastic mortality), self-thinning mortality, and the aforementioned mortality due to carbon starvation. In addition to mortality, biomass removal in 3D-CMCC-FEM results from forest management practices, such as thinning and final harvest (Collalti et al., 2018; Dalmonech et al., 2022; Testolin et al., 2023). The required model input data include stand age, average DBH (Diameter at Breast Height), stand density, and tree height (Collalti et al., 2014). The soil compartment is represented using one single bucket layer, in which the available soil water (ASW, in mm) is updated every day considering the water inflows (precipitation and, if provided, irrigation) and outflows (evapotranspiration, i.e., the sum of evaporation from the soil and transpiration of the canopy). The remaining water between these two opposite (in sign) fluxes that exceeds the site-specific soil water holding capacity is considered lost as runoff. For a full 3D-CMCC-FEM description, see: https://doi.org/10.32018/ForModLab-book-2024.

### 2.2 MEDFATE model

MEDFATE v.4.2.0 is an R-based modelling framework that allows the simulation of the function of forest ecosystems, with a specific emphasis on drought impacts under Mediterranean conditions (De Cáceres et al., 2021, 2023). MEDFATE calculates energy balance, photosynthesis, stomatal regulation, and plant transpiration of gas exchange separately at sub-daily steps. Like 3D-CMCC-FEM, MEDFATE also simulates photosynthesis at the leaf level using the biochemical model of Farquhar–von Caemmerer–Berry (Farquhar et al., 1980) for sunlit and shaded leaves (De Pury and Farquhar, 1997). MEDFATE can simulate plant hydraulics and stomatal regulation according to two different approaches: (a) steady-state plant hydraulics and optimality-based stomatal regulation (Sperry et al., 1998; Sperry et al., 2017); and (b) transient plant hydraulics including water compartments and empirical stomatal regulation (Sureau-ECOS; Ruffault et al., 2022). In this work, we took the second approach, i.e., Sureau-ECOS (Ruffault et al., 2022).

The hydraulic architecture of the Sureau-ECOS module comprises arbitrary soil layers, where the rhizosphere containing coarse and fine root biomass is calculated for each layer. The total root xylem conductance is determined by factors such as root length (limited by soil depth), weight, and distribution across the different layers. In addition, the resistance to water flow is dependent on two plant compartments (leaf and stem, each composed of symplasm and apoplasm). Overall, plant conductance is defined by the sum of resistances across the hydraulic network (i.e., soil, stem, and leaves), taking into account processes such as plant capacitance effects (i.e., the variation of symplasmic water reservoirs in the stem and leaves) and cavitation flows (i.e., water released to the streamflow from cavitated cells to non-cavitated cells during cavitation) (Hölttä et al., 2009). To withstand drought stress, adjacent conduits (tracheids or vessels) and/or living cells (e.g., parenchyma) release water to the xylem and may subsequently be refilled. In the event of embolization, cavitated xylem conduits release their water to the non-cavitated parts of the xylem, which then transfer it to adjacent compartments. Each element (roots, stem, leaves) of the hydraulic network has a vulnerability curve *kΨ*, that declines as water pressure becomes more negative. The xylem vulnerability curve is modelled using a sigmoid function, defined by the equation:

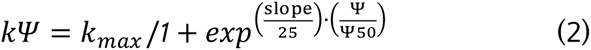

where *k_max_* is the maximum hydraulic conductance, *Ψ50* is the water potential corresponding to 50% of conductance, and “*slope*” is the slope of the curve at that point. The stem vulnerability curve can be used to determine the proportion of stem conductance loss (PLC_stem_) associated with vessel embolism. This embolism reduces overall tree transpiration and photosynthesis. Plant hydraulic failure and tree death can occur if the PLC_stem_ exceeds the 50% threshold.

Gas exchange in the Sureau-ECOS module depends on stomatal conductance (which depends on light, water availability, and air temperature) and leaf cuticular conductance, which changes with leaf temperature due to changes in the permeability of the epidermis. Stomatal regulation, unlike the 3D-CMCC-FEM, follows the Baldocchi (1994) approach, which allows coupling leaf photosynthesis with water losses. In addition, a multiplicative factor depending on leaf water potential is used to decrease stomatal conductance under drought conditions, following a sigmoidal function similar to stem vulnerability.

Soil water balance is computed daily. MEDFATE can consider an arbitrary number of soil layers with varying depths in which the water movement within the soil follows a dual-permeability model (Jarvis et al., 1991; Larsbo et al., 2005). Soil water content (ΔV_soil_, in mm) is calculated taking into account variables such as infiltration, capillarity rise, deep drainage, saturation effect, evaporation from the soil surface, transpiration of the herbaceous plant, and woody plant water uptake. A full MEDFATE description is available at: https://emf-creaf.github.io/medfatebook/index.html.

### 2.3 Evaluation sites

Model evaluation was performed in three PROFOUND and FLUXNET Network European beech sites, i.e., Sorø (DK-Sor, Denmark), Hesse (FR-Hes, France), and Collelongo (IT-Col, Italy), in which we retrieved information on soil texture, soil depth, and stand inventory data of forest structure for model initialization (Collalti et al., 2016; Marconi et al., 2017; Reyer et al., 2020a,b; https://fluxnet.org/). These sites are equipped with the Eddy Covariance towers (EC; Pastorello et al., 2020) for long-term continuous monitoring of atmospheric carbon, water, and energy fluxes of the forests (Fig. 1). The DK-Sor site is located in the forest Lille Bogeskov on the island of Zealand in Denmark. FR-Hes is situated in the northeastern region of France and lies on the plain at the base of the Vosges Mountains. IT-Col (Selva Piana stand) is a permanent experimental plot installed in 1991 and situated in a mountainous area of the Abruzzo region, centre of Italy.

**Fig 1.**
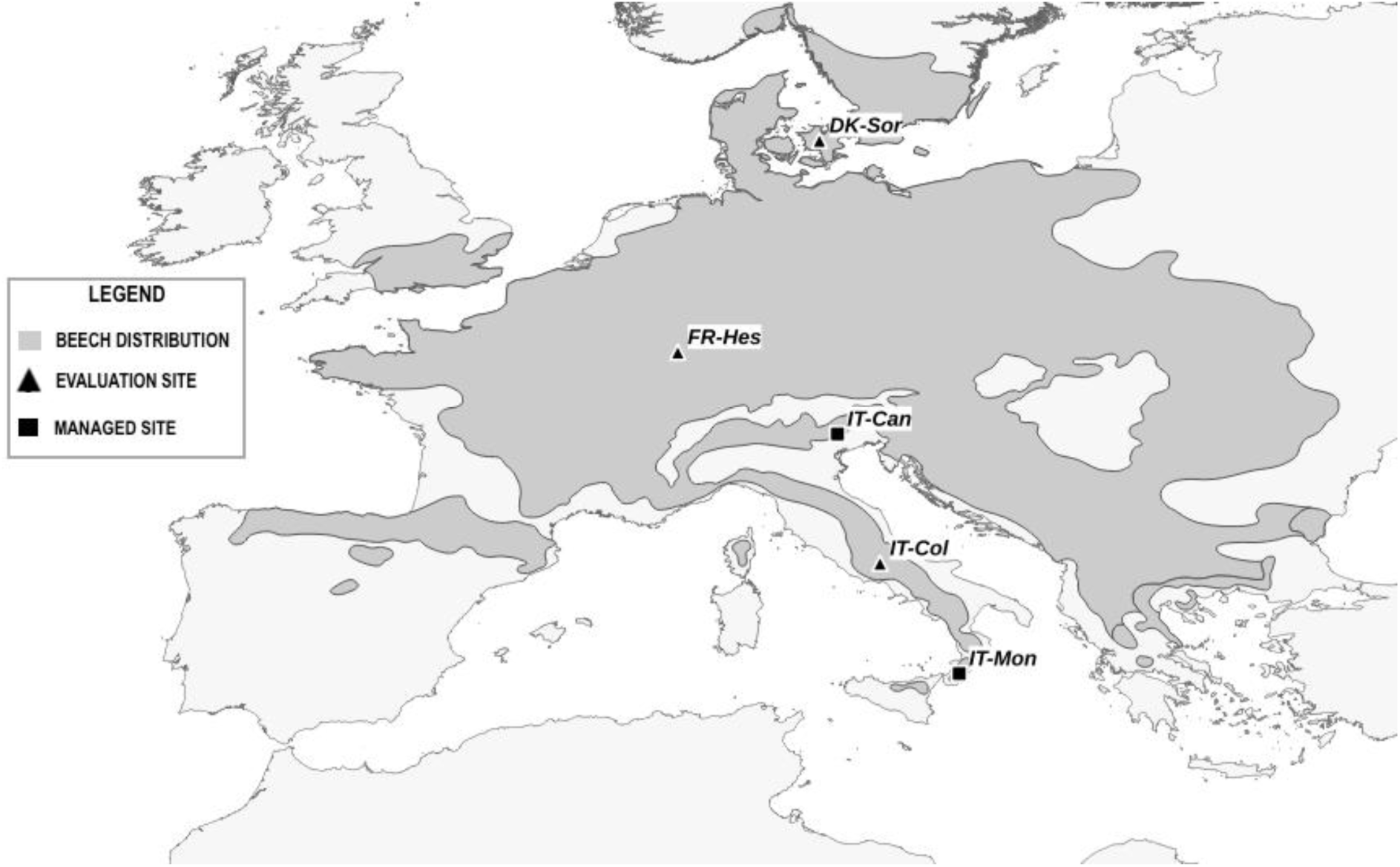
Map of the study sites. Triangles represent sites for validating fluxes, while the squares represent sites for management investigation.

The pedological characterization of soils exhibits distinct variations across the studied sites. The soil at the DK-Sor site is predominantly classified as either Alfisols or Mollisols. The FR-Hes site showcases an intermediary nature, displaying characteristics akin to both luvisols and stannic luvisols. At the IT-Col site, the prevailing soil type is identified as Humic alisols, according to the USDA soil classification system. Full details of these sites are reported in Table 1.

**Table 1.** Characteristics of the study sites. The age of the stands refers to 2010. The mean annual temperature (MAT) and mean annual precipitation (MAP) for DK-Sor, FR-Hes and IT-Col refer to the period evaluated (i.e., 2006-2010 for the Sorø and Collelongo site and 2014-2018 for the Hesse site) while for Cansiglio and Mongiana from 2010 to 2022. The sum of precipitation in summer refers to June (J), July (J) and August (A) for the same period.

| SITE DESCRIPTION |  |  |  |  |  |
| --- | --- | --- | --- | --- | --- |
| Evaluation sites |  |  |  | Test sites |  |
| Variable | DK-Sor | FR-Hes | IT-Col | Cansiglio | Mongiana |
| Coordinates (WGS84) | 55°49'N,<br>11°64' E | 48°66'N,<br>7°08'E | 41°85'N,<br>13°59'E | 46°02'N,<br>12°22'E | 38°29'N,<br>16°14'E |
| Country | Denmark | France | Italy | Italy | Italy |
| Altitude (m a.s.l.) | 40 | 305 | 1500 | 1300 | 1300 |
| Area (ha) | 1 | 1 | 1 | 27 | 27 |
| MAT (°C) | 8.52 | 10.27 | 6.95 | 6.44 | 11.01 |
| MAP (mm) | 818 | 853 | 1075 | 2219 | 1701 |
| Slope (%) | - | 5 | 35 | 12 | 10 |
| Aspect (°) | 0 | 0 | 252 | 135 | 135 |
| Stand age (yr) | 90 | 45 | 118 | 120 | 90 |
| Summer prec (J-J-A)(mm) | 292 | 205 | 120 | 493 | 141 |

The variables accounted for in the evaluation were obtained from the Fluxdata website (http://fluxnet.fluxdata.org/data/fluxnet2015-dataset/) from the FLUXNET2015 database (Pastorello et al., 2020). The variables considered are the daily GPP, estimated from Net Ecosystem Exchange (NEE) measurements and quality checked using the constant USTAR turbulence correction according to Papale et al. (2006) and the Latent Heat flux (LE) with energy balance closure correction (i.e., ‘LE_CORR’) (Pastorello et al., 2020).

### 2.4 Study sites

The two target sites considered in this study are Cansiglio and Mongiana Forests (Fig. 1) (De Cinti et al., 2016). Each site consists of nine long-term monitored plots of differently managed beech stands, with a spatial extension for each area above 3 hectares, for about 27 hectares of the experimental area. Three different silvicultural treatments were applied (see Fig. S1-S2). For each site, three of the nine plots considered were left unmanaged (i.e., no cutting and leaving the stands to natural development), defined as ‘Control’ plots, three plots were managed following the historical shelterwood system (‘Traditional’), and three with innovative cutting (‘Innovative’). In Cansiglio, considering the developmental stage of the stand was an establishment cut to open growing space in the canopy for the establishment of regeneration. The ‘Innovative’ cutting consisted of selecting a non-fixed number of scattered, well-shaped trees (the ‘candidate trees’) and a thinning of neighbouring competitors to reduce competition and promote better growth. In Mongiana, ‘Traditional’ silvicultural treatment was the first preparatory cut to increase the vitality and health of the intended residual trees in the stand. The ‘Innovative’ option was the identification of 45-50 as ‘candidate trees’ per hectare and removing only direct competitors.

The Cansiglio site is situated in a mountainous area in the Veneto region, northern Italy. Mongiana site is located in a mountainous area in the Calabria region of southern Italy. The latter shows higher mean annual temperature (MAT, C°) and lower mean annual precipitation (MAP, mm year^-1^) (i.e., drier conditions) than the Cansiglio site located at higher latitudes (Table 1). Data on forest structure and soil texture were collected during the field campaigns conducted in 2011 and 2019 (Cansiglio) and in 2012 and 2019 (Mongiana). At the Cansiglio site, soils are identified as Haplic luvisols, whereas at Mongiana, the predominant soil classifications consist of Inceptisols and Entisols, according to the USDA soil classification system. The variables analyzed in these sites, like in the evaluation sites, were GPP and LE. A summary of these sites is reported in Table 1.

### 2.5 Meteorological data

For the evaluation sites (i.e., DK-Sor, FR-Hes, IT-Col) observed meteorological data were retrieved from the harmonized PROFOUND database (Reyer et al., 2020a, b) and FLUXNET2015 database (Pastorello et al., 2020; https://data.icos-cp.eu/).

For the Mongiana and Cansiglio sites, meteorological data for 2010-2022 were obtained at daily temporal resolution from the relevant region’s Regional Environmental Protection Italian Agencies (ARPAs), which are responsible for monitoring climate variables with weather stations. The choice of thermo-pluviometric weather station was based on the minimum distance from the study area (between 2 km and 9 km away from the study sites, respectively) and on the data availability and integration with other weather stations in the proximity, representing the best available and obtainable meteorological observed data for these sites. The Bagnouls–Gaussen graph (Fig. S3) shows the mean monthly precipitation (mm) and air temperature (°C) recorded for every station inside the catchment.

Climate scenarios used as inputs for the two models at the Cansiglio and Mongiana sites were from the COSMO-CLM simulation at a spatial resolution of approximately 2.2 km over Italy (Raffa et al., 2023).

The daily variables considered for 3D-CMCC-FEM were mean solar radiation (MJ m^-2^ day^-1^), maximum and minimum air temperature (°C), precipitation (mm day^-1^), and the mean relative air humidity (%). In contrast, the MEDFATE model uses mean solar radiation, maximum and minimum air temperature, precipitation, the daily maximum and minimum relative air humidity, and wind speed (m s^-1^).

### 2.6 Modelling set-up

A set of parameters specific for *Fagus sylvatica* L. was provided as input to the model 3D-CMCC-FEM as described in Collalti et al. (2023) while for MEDFATE as in De Cáceres et al. (2023). To remove any confounding factors related to parameterization, the parameters related to photosynthesis and stomatal conductance were kept constant between the two models (see Table 2). A complete list of parameters and their values for both models as adopted in the present study can be found in supplementary materials (Table S2).

**Table 2.**
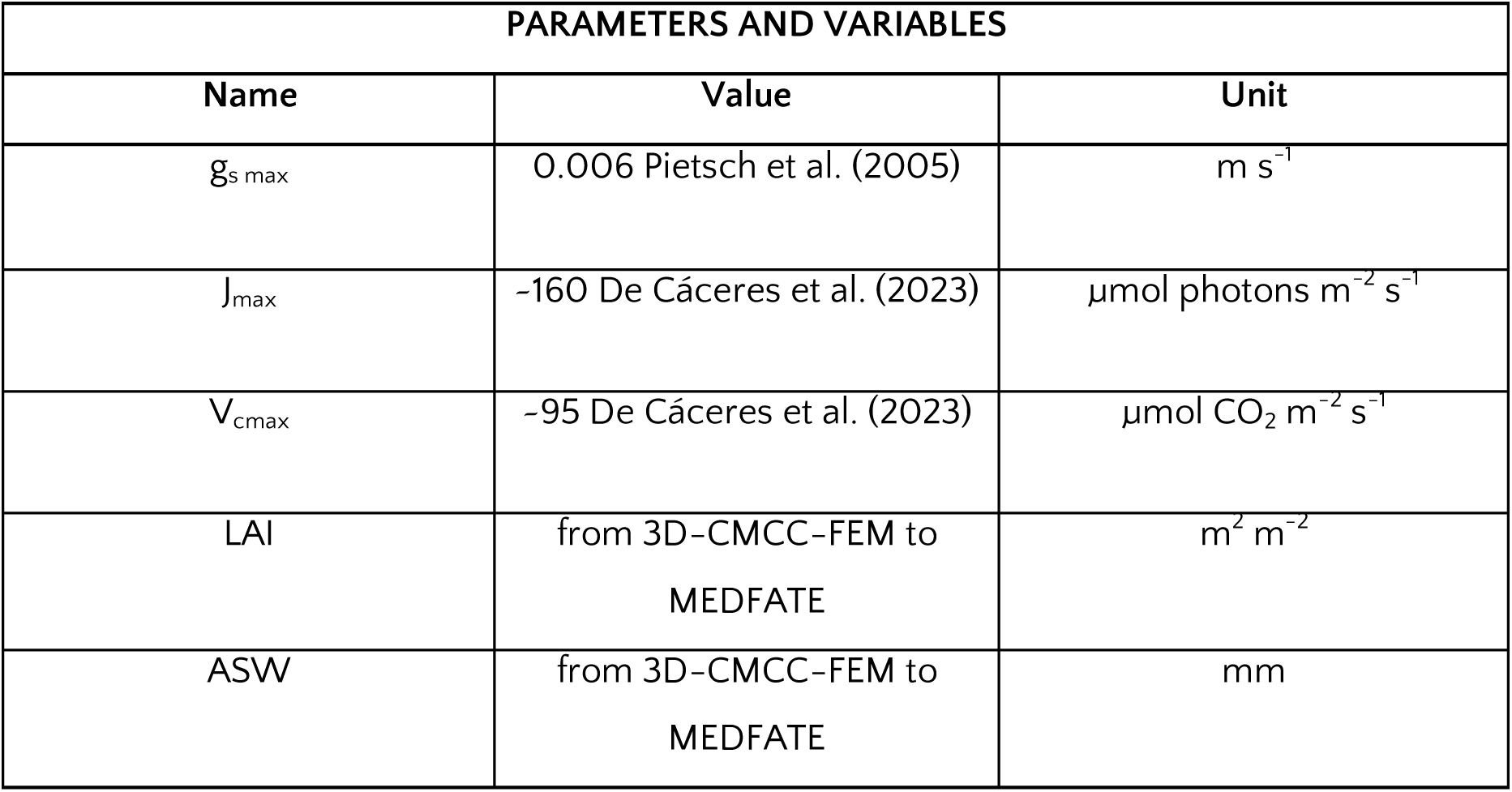
Parameters and variables set for both models during the simulations.

We then used the LAI and Available Soil Water (AWS) values obtained from the 3D CMCC-FEM outputs as input for running simulations with the MEDFATE model given that the model function used here does not prognostically simulate LAI and Available Soil Water (ASW). Precisely, here we used MEDFATE to simulate C and H_2_O fluxes only while considering plant hydraulics (De Cáceres et al., 2021), from the forest structure predicted, in terms of LAI, by 3D-CMCC-FEM. For MEDFATE water balance, LAI values determine the competition for light and also drive the competition for soil water, along with the root distribution across soil layers.

### 2.7 Model evaluation

To evaluate the performance of the two models across the different sites examined and under varying environmental conditions, we first assessed GPP and LE daily fluxes along a latitudinal gradient at sites equipped with EC towers. The reliability of the two models subsequently allowed us to coherently simulate fluxes at sites where EC towers were absent. Both models were run for five years on the evaluation sites, with the simulation period determined by the availability of observed data provided, as already mentioned, from the PROFOUND database, specifically, from 2006 to 2010 at DK-Sor and IT-Col sites while for FR-Hes starting from 2014 to 2018. The performance metrics of the results of the evaluation for each site for the GPP and LE variables were the coefficient of determination (R^2^), Root Mean Square Error (RMSE), and Mean Absolute Error (MAE).

### 2.8 Model application in managed sites

In the managed sites (i.e., Cansiglio and Mongiana), simulations were performed using Historical climate (‘Hist’) and, to analyse models’ sensitivities to climate change, under two Representative Concentration Pathways 4.5 and 8.5 (‘Moderate’ and ‘Hot Climate’), respectively. The ‘Hist’ climate was used to run simulations at the Cansiglio site from 2011 to 2022 and the Mongiana site from 2012 to 2022. In contrast, simulations using RCP4.5 and RCP8.5 climate ran accounting for the same period, that is, eleven years for the Cansiglio site and ten years for the Mongiana site, but considering the last years of the climate change scenarios (i.e., 2059-2070 and 2060-2070, respectively) to create harsher temperature and precipitation conditions, but with an increased atmospheric CO_2_ concentration (in μmol mol^-1^).

For each of the nine sampled areas, in the Cansiglio and Mongiana sites, we considered a representative area of one hectare for each type of plot: ‘Control’, ‘Traditional’, and ‘Innovative’. At the beginning of the simulations, each site thus included a total of 9 plots, each one hectare in size—comprising three ‘Control’ plots, three ‘Traditional’ plots, and three ‘Innovative’ plots. This setup resulted in a total of nine hectares being simulated per site where the model 3D-CMCC-FEM removed a certain percentage of the Basal Area (BA) according to the LIFE-ManFor project (see Table S1). ‘Traditional’ and ‘Innovative’ cutting took place for the first time in 2012 (Cansiglio) and 2013 (Mongiana), respectively. Following preliminary results, since the Mongiana site experienced a lighter thinning intensity compared to the Cansiglio site (refer to Table S1), consequently, for the Mongiana site, we considered an alternative management option involving the removal of 40% of the BA. This was done to evaluate whether a more intensive management approach (‘SM’) could have influenced models’ results on GPP and LE fluxes related to the reduction in competition and enhanced water availability.

## 3. Results

### 3.1 Model evaluation

The GPP at DK-Sor, FR-Hes, and IT-Col sites estimated from EC and simulated by 3D-CMCC-FEM and MEDFATE are shown in Fig. 2. At the DK-Sor site, the 3D-CMCC-FEM simulates a mean daily GPP of 5.14 gC m⁻ ² day⁻ ¹, while MEDFATE 5.13 gC m⁻ ² day⁻ ¹; and EC 5.54 gC m⁻ ² day⁻ ¹; at the FR-Hes site, 3D-CMCC-FEM mean daily GPP of 6.18 gC m⁻ ² day⁻ ¹ compared to MEDFATE 4.82 gC m⁻ ² day⁻ ¹, and EC 4.99 gC m⁻ ² day⁻ ¹; lastly at the IT-Col site, 3D-CMCC-FEM mean daily GPP of 4.88 gC m⁻ ² day⁻ ¹ compared to MEDFATE 4.19 gC m⁻ ² day⁻ ¹; and EC 4.11 gC m⁻ ² day⁻ ¹. Additionally, at the DK-Sor site, the 3D-CMCC-FEM simulated a mean daily LE of 2.83 MJ m⁻ ² day⁻ ¹, while the MEDFATE simulated a mean value of 2.22 MJ m⁻ ² day⁻ ¹; and EC 3.19 MJ m⁻ ² day⁻ ¹; at the FR-Hes site, 3D-CMCC-FEM simulated a mean daily LE of 4.16 MJ m⁻ ² day⁻ ¹ compared to MEDFATE 3.01 MJ m⁻ ² day⁻ ¹; and EC 4.47 MJ m⁻ ² day⁻ ¹; in the end at the IT-Col site, 3D-CMCC-FEM simulated a mean daily LE of 2.02 MJ m⁻ ² day⁻ ¹ compared to MEDFATE 2.57 MJ m^-^² day⁻ ¹; while EC 3.93 MJ m⁻ ² day⁻ ¹. The GPP predicted by 3D-CMCC-FEM has shown higher values of R^2^ (0.92) at DK-Sor and the lowest value at FR-Hes site (R^2^ = 0.76) whilst a value of R^2^ = 0.83 at IT-Col site, respectively. For the MEDFATE model, the highest predicted GPP value of R^2^ (0.85) was at DK-Sor, the lowest (R^2^ = 0.68) at IT-Col, and at FR-Hes R^2^ = 0.76, the same showed for the 3D-CMCC-FEM model, respectively. Differently, the highest R^2^ (0.89) value for 3D-CMCC-FEM considering LE predicted vs. observed was at FR-Hes site and almost the same values for DK-Sor and IT-Col sites (R*^2^* = 0.85 and 0.84, respectively). MEDFATE, for predicted vs. observed LE variable, has shown the highest R^2^ (0.77) at IT-Col site, lower R^2^ (0.69) value at FR-Hes site and the lowest R^2^ (0.62) value at DK-Sor site, respectively. In general, both the Root Mean Square Error (RMSE) and Mean Absolute Error (MAE) values in all sites were reasonably low, falling within the ranges of 3.31 to 2.02 gC m⁻ ² day⁻ ¹ and 2.46 to 1.47 MJ m⁻ ² day⁻ ¹, for both models and for both the variables. In Fig. 3 and Table 3 the summary of the evaluation metrics performance results.

**Fig 2.**
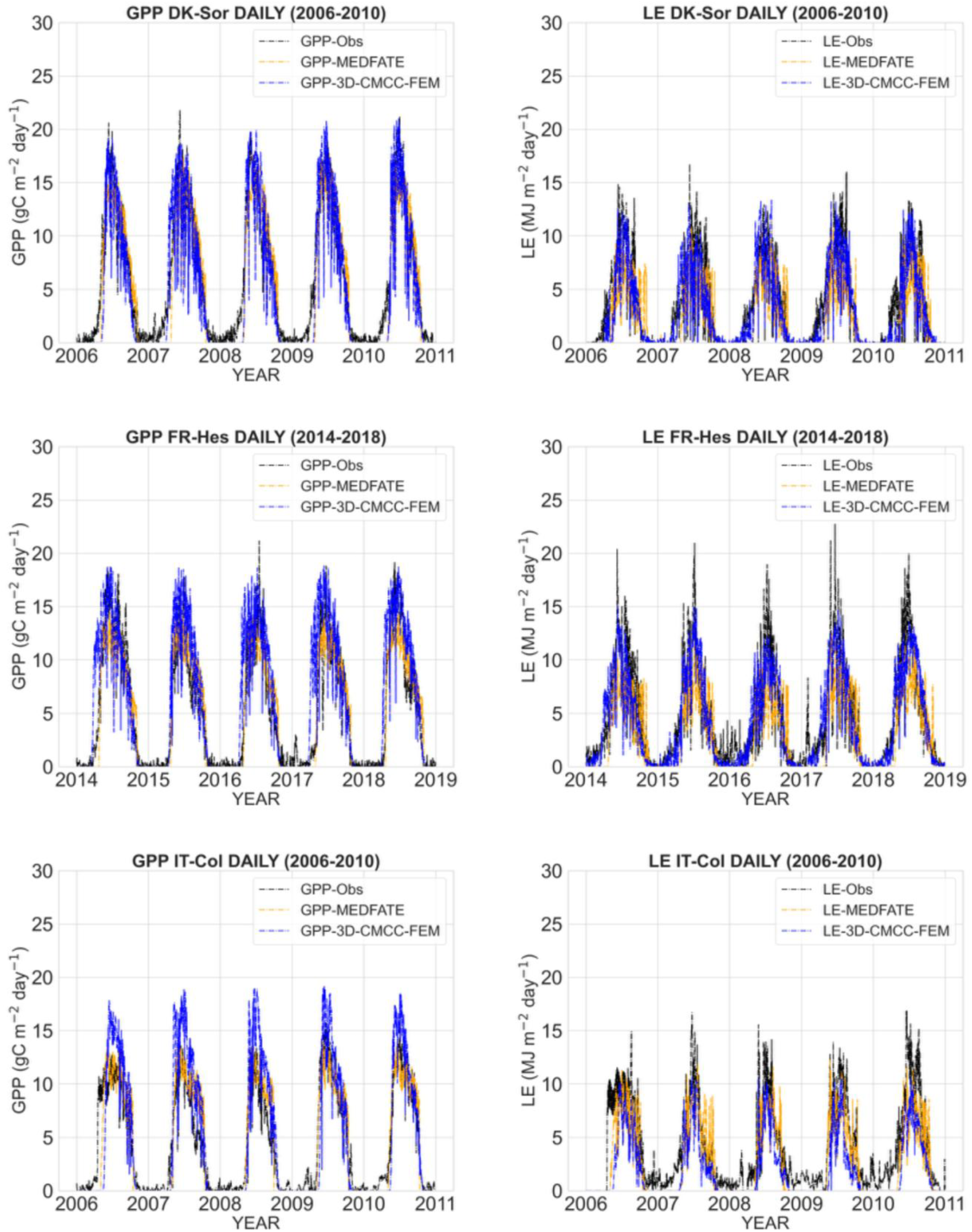
Daily mean variations of GPP (gC m⁻ ² day⁻ ¹) and LE (MJ m⁻ ² day⁻ ¹) estimated from the direct micrometeorological eddy covariance measurements (GPP*-*Obs and LE*-*Obs) and models’ simulation (GPP*-*3D-CMCC-FEM, LE*-*3D-CMCC-FEM and, GPP*-*MEDFATE, LE*-*MEDFATE) during the evaluation period at the DK-Sor, IT-Col and FR-Hes at the Beech forest in 2006-2010 and 2014-2018, respectively.

**Fig 3.**
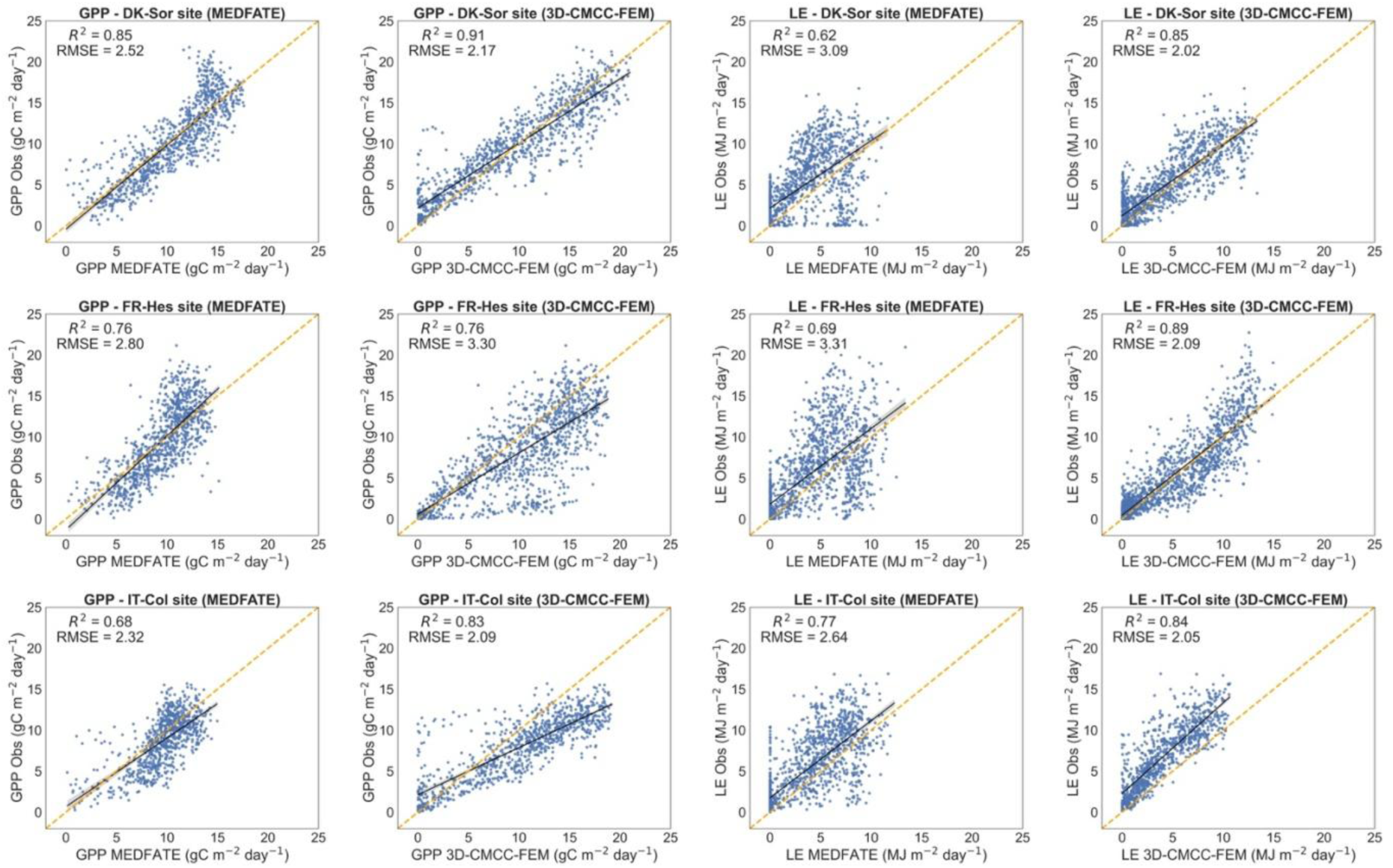
Scatter plots and linear regressions of GPP (gC m⁻ ² day⁻ ¹) and LE (MJ m⁻ ² day⁻ ¹) of the models versus the direct micrometeorological eddy covariance measurements (Obs) at the Sorø (DK-Sor; 2006-2010 period), Collelongo (IT-Col; 2006-2010 period) and Hesse (FR-Hes; 2014-2018 period).

**Table 3.** The correlation coefficient (R^2^), the Root Mean Square Error (RMSE) and the Mean Absolute Error (MAE) for, the GPP (gC m⁻ ^2^day⁻ ^1^) and LE (MJ m⁻ ^2^ day⁻ ¹) of the daily simulations at DK-Sor, IT-Col, and FR-Hes sites performed from both models 3D-CMCC-FEM and MEDFATE in the beech forest stands.

|  | 3D-CMCC-FEM |  |  |  |  |  | MEDFATE |  |  |  |  |  |
| --- | --- | --- | --- | --- | --- | --- | --- | --- | --- | --- | --- | --- |
|  | GPP |  |  | LE |  |  | GPP |  |  | LE |  |  |
|  | R<sup>2</sup> | RMSE | MAE | R<sup>2</sup> | RMSE | MAE | R<sup>2</sup> | RMSE | MAE | R<sup>2</sup> | RMSE | MAE |
| DK - Sor | 0.91 | 2.17 | 1.65 | 0.85 | 2.02 | 1.58 | 0.85 | 2.52 | 1.97 | 0.62 | 3.09 | 2.39 |
| FR - H | 0.76 | 3.30 | 2.46 | 0.89 | 2.09 | 1.47 | 0.76 | 2.80 | 2.21 | 0.69 | 3.31 | 2.37 |
| es |  |  |  |  |  |  |  |  |  |  |  |  |
| IT<br>-<br>C<br>ol | 0.<br>8<br>3 | 2.0<br>9 | 1.5<br>6 | 0.<br>8<br>4 | 2.0<br>5 | 1.<br>57 | 0.<br>6<br>8 | 2.3<br>2 | 1.<br>87 | 0<br>.7<br>7 | 2.6<br>4 | 1.<br>9<br>0 |

### 3.2 Simulation results at Cansiglio

Fig. 4 shows the simulation results using the 3D-CMCC-FEM and MEDFATE models in the Cansiglio site. For the 3D-CMCC-FEM, the ‘Control’ plot exhibited the lowest GPP values under ‘Hist’ climate conditions, averaging 1681 gC m⁻ ² year⁻ ¹. These values increased slightly to 1982 gC m⁻ ² year⁻ ¹ under the RCP4.5 climate and further rose to 2204 gC m⁻ ² year⁻ ¹ under the RCP8.5 climate. Similarly, for plots managed with ‘Traditional’ methods, the trends were consistent with the ‘Control’ plot, showing average GPP values of 1603, 1942, and 2141 gC m⁻ ² year⁻ ¹ under ‘Hist’, RCP4.5 and RCP8.5 climate, respectively. However, ‘Innovative’ management showed lower GPP fluxes across all three climate scenarios, with average values of 1534, 1882, and 2075 gC m⁻ ² year⁻ ¹ under ‘Hist’ RCP4.5 and RCP8.5 climate, respectively. The MEDFATE model showed higher mean absolute GPP increases than the 3D-CMCC-FEM model under RCP4.5 and RCP8.5 climates, respectively. Under the ‘Hist’ climate and all treatments, the mean GPP values were about 1638 gC m⁻ ² year⁻ ¹, whereas under the RCP4.5 climate, they rose to 2516 gC m⁻ ² year⁻ ¹ and 2995 gC m⁻ ² year⁻ ¹ under RCP8.5 climate. Analyzing in Fig. 4 the trends of LE for the 3D-CMCC-FEM model, these trends closely follow those of GPP concerning management treatments. The 3D-CMCC-FEM LE values for the ‘Control’ plots, similar to GPP, were lowest for the ‘Hist’ climate with an average value over the simulation years of 1200 and 1501 MJ m⁻ ² year⁻ ¹ for the RCP4.5 climate, and 1391 MJ m⁻ ² year⁻ ¹ for the RCP8.5 climate, respectively. The LE of the ‘Traditional’ management predicts values of 1129 in the ‘Hist’ climate, 1440 in the RCP4.5 climate, and 1338 MJ m⁻ ² yr⁻ ¹ in the RCP8.5 climate, respectively. For the ‘Innovative’ management, the mean LE values were 1121 in the ‘Hist’ climate, 14O3 for the RCP4.5 climate, and 1306 MJ m⁻ ² yr⁻ ¹ for the RCP8.5 climate, respectively. Similar to the GPP fluxes, the MEDFATE model simulated reductions in LE fluxes among the treatments and higher values across the climates. The mean LE value modelled in the ‘Hist’ climate, grouped by treatments (because of slight differences among managements), was about 920, 1419 in the RCP4.5 climate, and 1456 MJ m⁻ ² yr⁻ ¹ in the RCP8.5 climate, respectively (Fig. S12). MEDFATE simulated a stem xylem conductance loss of approximately 40% in the seventh, eighth, and twelfth years of simulation for the RCP8.5 climate scenario in the ‘Control’ plot. In contrast, this loss was predicted only in the seventh year for the managed plots. Conversely, near-zero or negligible stem embolism were simulated under the ‘Hist’ and RCP4.5 climate scenarios. The 3D-CMCC-FEM simulated higher values, albeit in a small percentage (i.e., between 8-10%) of NSC, increasing proportionally to the intensity of basal area removed, better observable in the graph at the tree level.

**Fig 4.**
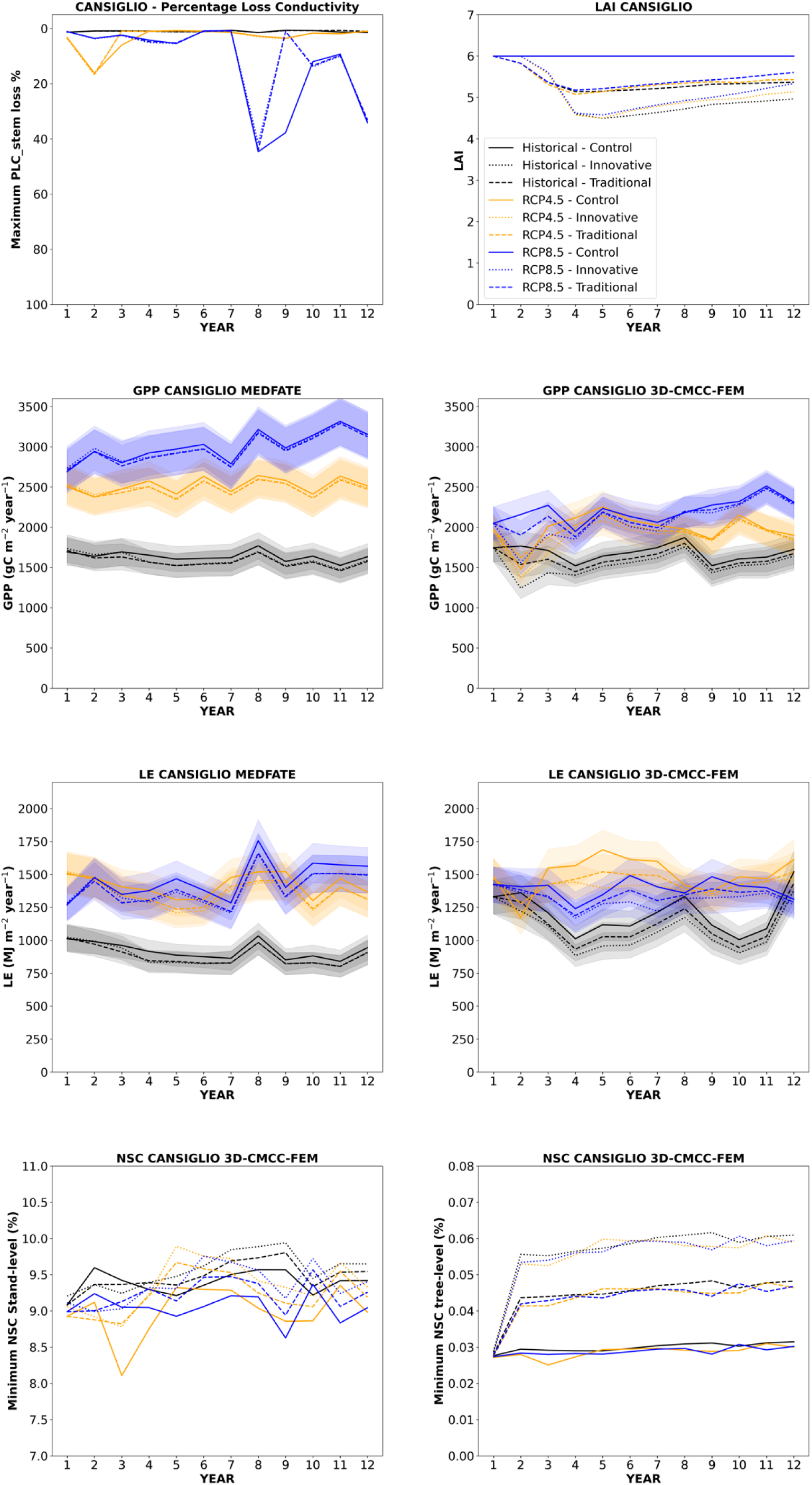
Comparative analysis between models output at the Cansiglio site. The top-left panel displays the PLC_stem_ as modelled by MEDFATE, while the top-right panel shows the modelled LAI for 3D-CMCC-FEM (and used by MEDFATE). The middle-up panels (left and right) present annual GPP (gC m⁻ ² year⁻ ¹) as modelled by the MEDFATE and 3D-CMCC-FEM, respectively. The middle-down panels (left and right) depict annual LE (MJ m⁻ ² yr⁻ ¹) modelled by the MEDFATE and 3D-CMCC-FEM, respectively. The bottom panels (left and right) depict the annual minimum of NSC concentration (%) at the stand and tree level, respectively, as modelled by the 3D-CMCC-FEM. Different plot management strategies are represented by distinct line styles: solid lines for ‘Control’ plots (’no management’), dotted lines for ‘Innovative’ plots, and dashed lines for ‘Traditional’ plots (Shelterwood). Climate scenarios are indicated by line colours: black for ‘Hist’ climate data (2010-2022), orange and blue for RCP4.5 and RCP8.5 climate (2059-2070), respectively.

### 3.3 Simulation results at Mongiana

Simulation results at the (drier) Mongiana site well depicted the differences with the rainy Cansiglio site (Fig. 5). The 3D-CMCC-FEM model showed no significant differences in the mean values of GPP among various management interventions under ‘Hist’ climate conditions, with a mean value of 2151 gC m⁻ ² yr⁻ ¹. Compared to the Cansiglio site, Mongiana exhibited lower average GPP values. Under RCP4.5 climate conditions, the GPP for the ‘Control’ plot was 1864 gC m⁻ ² yr⁻ ¹. In contrast, under the current climate, the ‘Traditional’ and ‘Innovative’ management interventions yielded higher average GPP values of 2115 gC m⁻ ² yr⁻ ¹ and 2086 gC m⁻ ² yr⁻ ¹. The GPP values under the more intensive management (‘SM’) and RCP4.5 climate decreased even further than those of the ‘Control’ plot, with an average value of 165O gC m⁻ ² yr⁻ ¹. Under the RCP8.5, no differences in GPP were observed among management strategies, with values of about 1525 gC m⁻ ² yr⁻ ¹. Moreover, under the RCP8.5, the ‘Control’ plot experienced complete mortallty after five years of simulations. The MEDFATE model predicted slightly higher average values of GPP in the ‘Control’ plots (2099 gC m⁻ ² yr⁻ ¹) compared to the managed plots (2087 gC m⁻ ² yr⁻ ¹, encompassing both ‘Traditional’ and ‘Innovative’ of two management strategies), with no significant differences observed among the management strategies and under ‘Hist’ climate. Under the RCP4.5 and RCP8.5, the GPP values were 1608 gC m⁻ ² yr⁻ ¹ and 1935 gC m⁻ ² yr⁻ ¹, respectively (Fig. S11). The PLC_stem_ graph in Fig. 5 indicated very high xylem embolism levels (i.e., reaching 100% every year) under RCP4.5 and RCP8.5 already in the first year of simulations. A pronounced embolism event was observed under the ‘Hist’ climate in 2017, 2018, 2019, and 2022 in a 30-45% range for the ‘Control’ plots, while the managed plots experienced a maximum embolism of approximately 40% in 2017. Conversely, the 3D-CMCC-FEM model did not report any significant differences between managed and unmanaged plots for the LE. The average LE value for the ‘Hist’ climate was 1796 MJ m⁻ ² yr⁻ ¹, which decreased to 1220 MJ m⁻ ² yr⁻ ¹ under the RCP4.5 and 1190 MJ m⁻ ² yr⁻ ¹ under the RCP8.5 in managed plots. As previously described, the ‘Control’ plot under the RCP8.5 experienced mortallty in the sixth year of simulation. Similarly to the previously described GPP fluxes, the MEDFATE model reported a slight difference in LE fluxes between the ‘Control’ plot under historical climate conditions (1623 MJ m⁻ ² yr⁻ ¹) and the managed plots (1603 MJ m⁻ ² yr⁻ ¹). For the RCP4.5 and RCP8.5, the LE values were 1100 MJ m⁻ ² yr⁻ ¹ and 1089 MJ m⁻ ² yr⁻ ¹, respectively (Fig. S12). The LE values of the ‘Control’ plots are not reported either for RCP4.5 or for RCP8.5 because of the mortality experienced for the simulation years.

**Fig 5.**
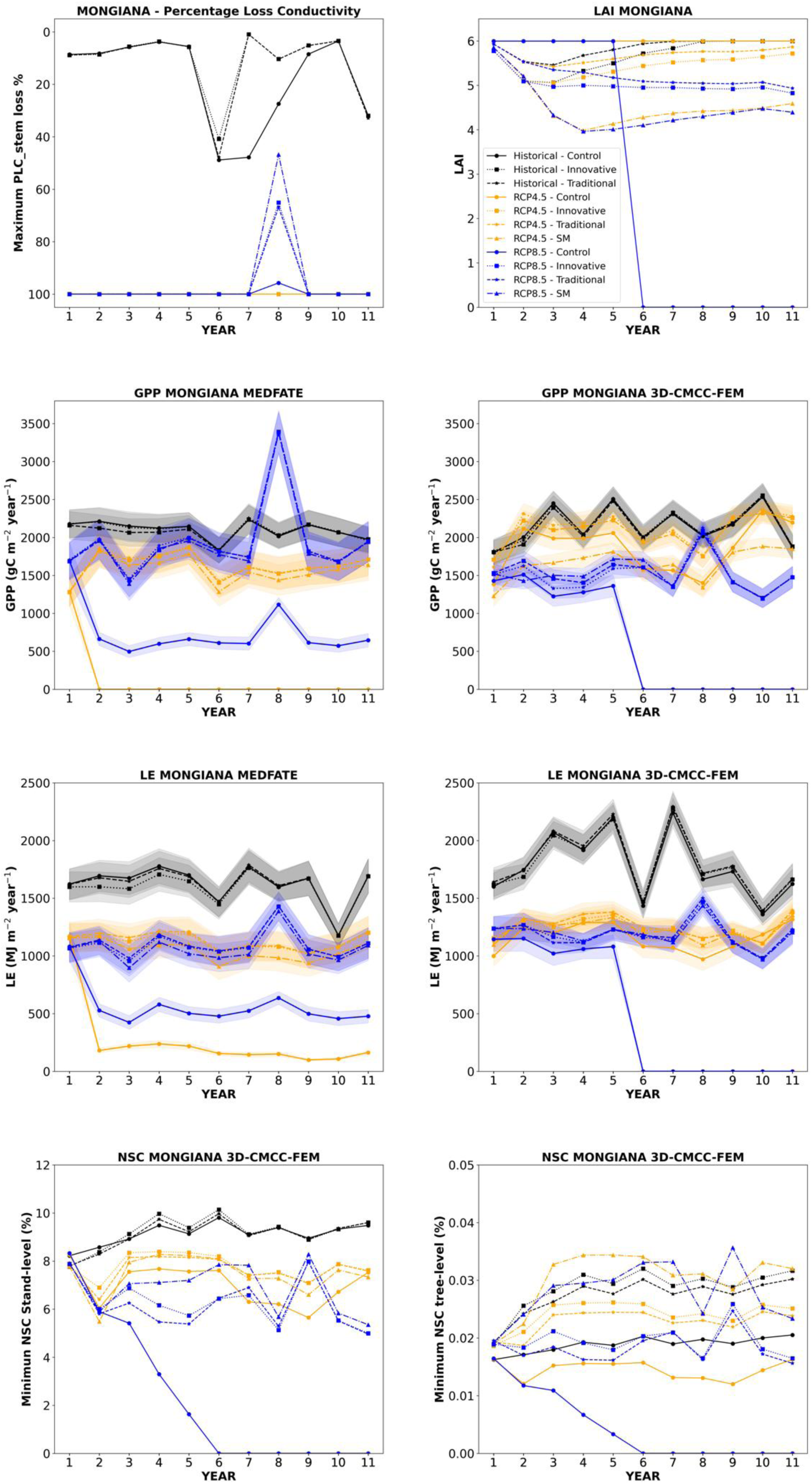
Comparative analysis between models output at the Mongiana site. The top-left panel displays the percent loss of PLC_stem_as modelled by MEDFATE while the top-right panel shows the modelled LAI for 3D-CMCC-FEM (and used by MEDFATE). The middle-up panels (left and right) present annual GPP (gC m⁻ ² year⁻ ¹) as modelled by the MEDFATE and 3D-CMCC-FEM, respectively. The middle-down panels (left and right) depict annual LE (MJ m⁻ ² yr⁻ ¹) modelled by the MEDFATE and 3D-CMCC-FEM, respectively. The bottom panels (left and right) depict the annual minimum of NSC concentration (%) at the stand and tree level, respectively, as modelled by the 3D-CMCC-FEM. Different plot management strategies are represented by distinct line styles: solid lines with circles for ‘Control’ plots (‘no management’), dotted lines with squares for ‘Innovative’ plots, dashed lines with stars for ‘Traditional’ plots (Shelterwood) and dash-dotted lines with triangles for ‘SM’ management. Climate scenarios are indicated by line colours: black for ‘Hist’ climate data (2010-2022), orange and blue for RCP4.5 and RCP8.5 climate (2060-2070), respectively.

## 4. Discussions

First, this study evaluated the performances of two different process-based models in simulating diverse beech stands across Europe, starting to the north of Europe and moving towards the south under different environmental conditions. Secondly, the study has focused on the models’ sensitivity and the relative impacts of different management options and different climatic conditions in two independent beech forest stands in the north and south of the Italian peninsula.

### 4.1 Model evaluation

To assess the models’ accuracy in predicting C and H_2_O fluxes, we compared daily GPP and LE data obtained from the EC towers. Both models predicted daily GPP and LE accurately and ensured a good range of general applicability of both models (Kramer et al., 2002; Verbeeck et al., 2008). The 3D-CMCC-FEM model seems to slightly overestimate GPP daily values along latitudinal gradients starting from the north (DK-Sor) to the south (IT-Col), as already found in Collalti et al. (2016). MEDFATE, in contrast, showed a slight overestimation of GPP only at IT-Col site. The LE predicted by 3D-CMCC-FEM is more accurate than MEDFATE prediction for DK-Sor and FR-Hes sites but not in IT-Col site in which 3D-CMCC-FEM has shown to underestimate compared to the observed EC values. For MEDFATE the underestimation of LE was observed in all the evaluation sites.

The spread observed for the GPP and LE fluxes between the two models may be attributed to the different assumptions that govern stomatal regulation since both models use the Farquhar-von Caemmerer-Berry biochemical model to calculate photosynthesis. The over or underestimation of the flows estimated by the models both for GPP and the LE compared to the data observed from the EC towers can be attributed either to the presence of the understory (although commonly sporadic in mature beech stands), which was not considered in the simulations by both models and to errors on daily measurement by EC technique (Loescher et al., 2006) or because a not perfect fit in the modeled seasonality (i.e., the begin and the end of the growing season) (Richardson et al., 2010). However, the overall leaf phenological pattern of the European beech in these sites is well represented by the two models in almost all of the years according to EC data as shown in supplementary materials (see Fig. S4, S5, S6, S7, S8, S9, and Fig S10). It is important to note that we did not specifically callbrate the models’ parameters at each site separately. Instead, both models were parameterized using existing values taken from the literature, therefore with one single set of parameter values for all sites.

### 4.2 Climate change and forest management at the Cansiglio and Mongiana site

The pre-Alpine site of Cansiglio showed slight differences in the fluxes (i.e., GPP and LE) between the three different management practices and the three climate scenarios (i.e., no climate change, RCP4.5. and RCP8.5) used (Fig. S11–S12). Future climate is expected to be higher temperature if compared to the historical one, with MAT higher of about 3.93°C under RCP4.5 and 4.95°C under RCP8.5 for the Cansiglio site and 4.52°C under RCP4.5 and 5.42°C under RCP8.5 at the Mongiana site. Similarly, MAP is expected to be 510 mm lower under RCP4.5 and 602 mm under RCP8.5 at the Cansiglio site, respectively, while 902 mm lower under RCP4.5 and 914 mm under RCP8.5 at the Mongiana site, respectively.

Regarding management, the response of the 3D-CMCC-FEM to the removal of a percentage of the basal area from the stand led to a decrease in GPP in the ‘Traditional’ cutting and an even greater extent, in the ‘Innovative’ cutting compared to the ‘Control’ (i.e., no management). Similarly to 3D-CMCC-FEM, the MEDFATE model simulates slight differences influxes amount (e.g., lower values for ‘Traditional’ and ‘Innovative’ cutting than ‘Control’ plots) between the management regimes in the plots. These results align with those of Guillemot et al. (2014), who observed a slight decrease in GPP in managed compared to unmanaged temperate beech forests in France under different thinning regimes. However, differences were observed in both models under the three different climates used in the simulations. GPP increased from the ‘Hist’ climate to RCP4.5 and reached the maximum for RCP8.5, respectively. This suggests a plastic response (e.g., photosynthesis and stomatal response) of the stands, as simulated by models, to harsher conditions, indicating, potentially, a high drought acclimation capacity (Petrik et al., 2022) and increased GPP because of the early budburst, and a prolonged vegetative season (Peano et al., 2019), and the so-called ‘atmospheric CO_2_ fertilization’ effect as also found by de Wergifosse et al. (2022) and Reyer et al. (2013), especially in sites with no apparent water limitation both under current and projected future climate conditions. The anisohydric behavior of *Fagus sylvatica* L. results in prolonged stomatal opening relative to isohydric species, although Puchi et al. (2024) recently found large variability in European beech responses, maintaining prolonged photosynthetic activity. Still, this response is modulated by summer precipitation and the availability of soil water storage (Leuschner et al., 2021; Baudis et al., 2015). However, for high-altitude stands, growth could be negatively affected under warmer conditions, as suggested by Chmura et al. (2024). The LE results for the 3D-CMCC-FEM showed lower values over the simulation period for managed stands than unmanaged ones showing lesser sensitivity to forest management if compared to MEDFATE. Yet, under the RCP4.5, the LE values were higher compared to both the ‘Hist’ climate and the RCP8.5 one due to greater annual cumulative precipitation than the RCP8.5 and higher, on average, temperatures than the ‘Hist’ scenario (Fig. S12).

Conversely, the MEDFATE model was shown to be more sensitive to climate, with a clearer distinction between the ‘Hist’ climate, the RCP4.5 and RCP8.5 climates, with higher and nearly equal values in the harsher conditions (i.e., RCP4.5 and RCP8.5 climates), with slight differences in the management treatments as obtained by 3D-CMCC-FEM.

The Non-Structural Carbon (NSC) amount showed the highest values in ‘Innovative’ plots, followed by ‘Traditional’ plots, and the lowest values in ‘Control’ plots, suggesting a benefit in carbon stock accumulation with more carbon going for carbon biomass and less for reserve-replenishment for these stands under management interventions. Nevertheless, NSC levels remain nearly the same for the three climate scenarios throughout all the simulation years. It is important to note that MEDFATE simulated an initial loss of stem conductance under the climate scenarios, indicating a premature onset of water stress for the stand. Although in RCP4.5 this is negligible, in RCP8.5 PLC_stem_ values reach a maximum xylem cavitation value of about 40% in the eighth year of simulation for managed plots while for ‘Control’ plots in the eighth, ninth, and twelfth years, highlighting potential benefits of management to reduce drought stress because of less rain interception and canopy evaporation and transpiration (Giuggiola et al., 2018; Schmied et al., 2023).

The GPP at the southern Apennine site of Mongiana showed a decrease under RCP4.5 and RCP8.5 scenarios when simulated by the 3D-CMCC-FEM model as a result of harsher environmental conditions, as also resulted in the study by Yu et al. (2022), in which the productivity and then the growth of European beech in southern regions are expected to decrease as affected by more severe climate conditions such as decreased precipitation and increased air temperature (Tognetti et al., 2019). Indeed, the increase in air temperature, a reduction in soil water availability, and the rise in vapor pressure deficit (VPD) lead to earlier stomatal closure, increased mesophyll resistance, and elevated abscisic acid production (Kane and McAdam, 2023), all of which contribute to a decrease in the carbon assimilation rate (Priwitzer et al., 2014; Grossiord et al., 2020). Specifically, GPP is higher under ‘Hist’ climate conditions, decreases under the RCP4.5, and ultimately reaches even lower values under the RCP8.5. Under the RCP8.5 at the fifth year of simulation, the stand in the ‘Control’ plot is simulated to die due to carbon starvation. The annual decline in NSC (Fig. 5) due to an imbalance between carbon uptake (photosynthesis) and the demands for growth and respiration suggests that the trees are unable to replenish their carbon reserves. The depletion of NSC reserves may ultimately disrupt processes such as osmoregulation and phenology (Martínez-Vilalta et al., 2016), potentially leading to stand defoliation and/or mortality. The management options did not show changes in GPP under the ‘Hist’ climate. However, the increase of GPP was observed under the RCP4.5 in the plots where ‘Innovative’ and ‘Traditional’ cutting occurred, although no differences were observed between them. For instance, the same increase in GPP was reported by Fibbi et al. (2019) for other European beech forests under climate change scenarios in Italy. The thinning reduces the leaf area and then the LAI and increases the soil water availability, which positively influences stomatal conductance and carbon assimilation, providing an acclimation mechanism to drought during periods of water scarcity (Lüttschwager and Jochheim, 2020; Diaconu et al., 2017).

In contrast, the more intense cutting exhibited even lower GPP values than the ‘Control’ plots. This is likely due to the overly intense thinning, which contrasts the microclimate effects within this forest stand, reducing the potential to offset climate warming at the local scale (Rita et al., 2021). Heavy thinning, on the other hand, can increase light penetration, soil evaporation, and wind speed, thereby heightening tree sensitivity to vapor pressure deficit under dry conditions (Schmied et al., 2023; Simonin et al., 2007). LE decreased with the decrease in precipitation under the RCP4.5 and RCP8.5 climate scenarios compared to the ‘Hist’ climate. There were no significant differences in LE among the various management regimes. For the MEDFATE model, negligible or no differences in GPP were observed under all the climates among various management options. Although the GPP values estimated by the MEDFATE model under the RCP4.5 and RCP8.5 are similar to those obtained from the 3D-CMCC-FEM model, a closer analysis of the daily outputs (data not shown) reveals that trees photosynthesize until the end of July, after which they experience significant embolism (i.e., maximum value of 100%), as indicated by the PLC_stem_ graph, indicating that the decrease in precipitation led to summer soil moisture depletion and lethal drought stress levels.

Furthermore, the ‘Control’ plots experienced mortallty even before reaching the summer period. In recent decades, prolonged drought stress in Mediterranean mountain regions has significantly reduced the productivity of beech forests, resulting in a decline in Basal Area Increment (BAI) and overall growth (Piovesan et al., 2008). It is also important to note that under ‘Hist’ climate conditions, the MEDFATE model indicated a stem embolization loss ranging from approximately 10% to 45% during the drought period (i.e., 2018-2020) in Europe as also highlighted in other study (Italiano et al., 2024; Thom et al., 2023; Lombardi et al., 2023). The embolization was more pronounced and long-lasting in the ‘Control’ plots than the managed ones. The same trends were obtained for LE.

### 4.3 Uncertainties and factors influencing forest carbon and water dynamics

Although there is scientific evidence of the positive effects of CO_2_ fertilization effect on forest primary productivity, uncertainties remain regarding the long-term persistence of this positive feedback and the level at which this may saturate (Sperlich et al., 2020; Wang et al., 2020). Furthermore, the down-regulation of this fertilization effect on photosynthesis is influenced by interannual variations in meteorological parameters, as well as by interactions within the carbon and nitrogen cycles. These factors must be carefully assessed to improve the accuracy of flux projections under future climatic conditions (Zaehle et al., 2014). It is also worth highlighting that this study does not account for biotic disturbances such as pest outbreaks and diseases, nor for abiotic disturbances like extreme climatic events (e.g., heatwaves, late frosts, and wildfires). These factors could significantly alter carbon and water fluxes (Yu et al., 2022b), potentially depleting carbon reserves faster and reducing the capacity for carbon sequestration by these forest ecosystems as well as possibly necessitating the adoption of different management strategies (Langer and Bußkamp, 2023; Margalef-Marrase et al., 2020). Another critical factor is the depth of the root zone and soil, as well as its physico-chemical composition. For example, it has been found that two-thirds of fine roots in European beech are within the top 30 cm of soil, while coarse roots can extend beyond depths of 240 cm (Meier et al., 2017). Although findings by Gessler et al. (2021) indicate that, unlike oak forests, European beech forests cannot compensate for additional water uptake from deeper soil layers during drought periods, Brinkmann et al. (2018) reported contrasting results. In this study, we analyzed soil texture characteristics to a depth of 110 cm for Cansiglio and 40 cm for Mongiana. However, the limited understanding of deeper layers may not fully capture the entire soil water reservoir and its dynamics. Expanding knowledge of the deepest soil layers is essential to better understand root development and, consequently, improve water storage capacity and drought resilience in beech forests.

## 5. Conclusions

The two process-based models provide robust evidence for their application in estimating fluxes, consistent with long-term EC tower measurements in European beech forests. Despite the minimal parametrization effort to align the two models and the avoidance of single site-specific parameters, reliable results can still be obtained, as confirmed by the outputs from the Sorø, Hesse, and Collelongo sites. Regarding the sub-Alpine Cansiglio site, although water limitation does not significantly impact fluxes or the health of the forest under ‘Moderate’ climate conditions (RCP4.5), a potential concern is the embolization predicted by the MEDFATE model under the ‘Hot’ climate (RCP8.5) at this site, despite similar levels of precipitation. The high susceptibility of European beech forests at the southern Apennine site of Mongiana to more severe (i.e., hotter and drier) climatic conditions could lead to the collapse of this forest ecosystem, even with the application of management options to reduce competition. However, it is crucial that these seasonal droughts are not prolonged or intense enough to exceed the ecological limits of the European beech. To avoid that European beech forests may necessitate of strategic and specifically designed management planning at the single site level, including the ability to project (e.g., with forest models) and evaluate future forest conditions for better management schemes. However, the ability of these forests to survive or resist the impacts of climate change may not depend solely on density reduction interventions. Prioritizing the exploration of alternative sustainable management strategies to promote carbon sequestration in both above-ground biomass and soil is crucial for enhancing climate change mitigation efforts. Additionally, evaluating silvicultural plans such as the introduction of complementary species can improve the resilience of vulnerable European beech ecosystems. A modeling approach, similar to the one used in this study, offers a valuable tool for assessing these alternative strategies and refining forestry adaptive management practices. By integrating these approaches, we can strengthen the long-term sustainability of forests while preserving the ecological balance of vulnerable regions.

## CRediT authorship contribution statement

**Vincenzo Saponaro:** Conceptualization, Data curation, Formal analysis, Investigation, Methodology, Resources, Software, Visualization, Writing – original draft, Writing – review & editing. **Miquel De Càceres:** Conceptualization, Methodology, Software, Supervision, Writing – review & editing. **Daniela Dalmonech:** Conceptualization, Methodology, Software, Supervision, Writing – review & editing. **Ettore D’Andrea:** Resources, Methodology, Data curation, Writing – review & editing. **Elia Vangi:** Resources, Writing – review & editing. **Alessio Collalti:** Conceptualization, Methodology, Resources, Software, Supervision, Project administration, Writing – review & editing.

## Data availability

Data will be made available on request.

## Supporting information

https://drive.google.com/file/d/1kgLPmNZndxx0PmFVKHPs6YmR1clTPLSf/view?usp=share_link

## Acknowledgements

We would like to thank the University of Tuscia, the CNR ISAFOM of Perugia and the company ENI s.p.a. for the realization of this work. Special thanks to the Institute Research Center for Ecological and Forestry Applications (CREAF) of Barcelona that supported the research by the Spanish “Ministerio de Ciencia e Innovación” (MCIN/AEI/10.13039/501100011033) (grant agreement No. PID2021-126679OB-I00).

The authors sincerely thank the ARPA Veneto and Calabria for providing the meteorological data. The Fondazione Centro Euro-Mediterraneo sui Cambiamenti Climatici (CMCC) for providing climate change scenarios. The Cost-Action PROFOUND for providing both the stand ancillary data and the climate used in this work. D.D., E.V. and A.C. has been partially supported by MIUR Project (PRIN 2020) “Unravelling interactions between WATER and carbon cycles during drought and their impact on water resources and forest and grassland ecosySTEMs in the Mediterranean climate (WATERSTEM)” (Project number: 20202WF53Z), “WAFER” at CNR (Consiglio Nazionale delle Ricerche) and by PRIN 2020 (cod. 2020E52THS) - Research Projects of National Relevance funded by the Itallan Ministry of University and Research entitled: “Multi-scale observations to predict Forest response to pollution and climate change” (MULTIFOR, project number: 2020E52THS). A.C. acknowledge also funding by the project OptForEU Horizon Europe research and innovation programme under grant agreement No. 101060554. D.D. and A.C. also acknowledge the project funded under the National Recovery and Resilience Plan (NRRP), Mission 4 Component 2 Investment 1.4 - Call for tender No. 3138 of 16 December 2021, rectified by Decree n.3175 of 18 December 2021 of Italian Ministry of University and Research funded by the European Union – NextGenerationEU under award Number: Project code CN_00000033, Concession Decree No. 1034 of 17 June 2022 adopted by the Italian Ministry of University and Research, CUP B83C22002930006, Project title “National Biodiversity Future Centre - NBFC”. This work used eddy covariance data acquired and shared by the FLUXNET community, including these networks: AmeriFlux, AfriFlux, AsiaFlux, CarboAfrica, CarboEuropeIP, CarboItaly, CarboMont, ChinaFlux, Fluxnet-Canada, GreenGrass, ICOS, KoFlux, LBA, NECC, OzFlux-TERN, TCOS-Siberia, and USCCC. The 3D-CMCC-FEM model code is publicly available and can be found on the GitHub platform at: https://github.com/Forest-Modelling-Lab/3D-CMCC-FEM. The MEDFATE model code is publicly available and can be found on the GitHub platform at: https://github.com/emf-creaf/medfate.

