## Supplementary material for "Assessing the combined effects of forest management and climate change on carbon and water fluxes in European beech forests": https://drive.google.com/file/d/1kgLPmNZndxx0PmFVKHPs6YmR1clTPLSf/view?usp=share_link

Table S1. Mean percentage (%) of BA removed for each plot (P) in each site.

| MANAGEMENT DESCRIPTION | | | | | | | | | | | | | | | | | |
|---|---|---|---|---|---|---|---|---|---|---|---|---|---|---|---|---|---|
| **Cansiglio** | | | | | | | | | **Mongiana** | | | | | | | | |
| **Control** | | | **Traditional** | | | **Innovative** | | | **Control** | | | **Traditional** | | | **Innovative** | | |
| P1 | P2 | P3 | P1 | P2 | P3 | P1 | P2 | P3 | P1 | P2 | P3 | P1 | P2 | P3 | P1 | P2 | P3 |
| - | - | - | 33 | 33 | 33 | 47 | 47 | 47 | - | - | - | 16 | 16 | 16 | 21 | 21 | 21 |

Table S2. MEDFATE and 3D–CMCC–FEM model parameters

| MEDFATE parameters | | | |
|---|---|---|---|
| **Parameter Name** | **Value** | **Unit** | **Description** |
| SNFI23/34 (Spain) | 0.106 | $cm^2 \cdot cm^{-1} \cdot yr^{-1}$ | Average relative annual growth rates for tree taxa |
| SNFI23 (Spain) | 0.113 | $cm^2 \cdot cm^{-1} \cdot yr^{-1}$ | Average relative annual growth rates for tree taxa |
| SNFI34 (Spain) | 0.102 | $cm^2 \cdot cm^{-1} \cdot yr^{-1}$ | Average relative annual growth rates for tree taxa |
| SNFI23/34 (Catalonia) | 0.138 | $cm^2 \cdot cm^{-1} \cdot yr^{-1}$ | Average relative annual growth rates for tree taxa |
| SNFI23 (Catalonia) | 0.149 | $cm^2 \cdot cm^{-1} \cdot yr^{-1}$ | Average relative annual growth rates for tree taxa |
| SNFI34 (Catalonia) | 0.110 | $cm^2 \cdot cm^{-1} \cdot yr^{-1}$ | Average relative annual growth rates for tree taxa |
| GrowthForm | Tree | | Growth form: Either "Shrub", "Tree" or "Tree/Shrub" |
| LifeForm | Phanerophyte | | Raunkiaer life form |
| LeafShape | Broad | | Broad/Needle/Linear/Scale/Spines/Succulent |
| LeafSize | Medium | | Either "Small" (< 225 mm), "Medium" (> 225 mm & < 2025 mm) or "Large" (> 2025 mm) |
| PhenologyType | winter-deciduous | | Leaf phenology type |
| Hmed | 1600 | cm | Median plant height |
| Hmax | 3200 | cm | Maximum plant height |
| Z50 | | mm | Depth corresponding to 50% of fine roots |
| Z95 | 867 | mm | Depth corresponding to 95% of fine roots |
| fHDmin | 50 | | Minimum value of height-diameter ratio |
| fHDmax | 160 | | Maximum value of height-diameter ratio |

| | | | |
|---|---|---|---|
| a_ash | | | Allometric coefficient for shrub area as function of height |
| b_ash | | | Allometric coefficient for shrub area as function of height |
| a_bsh | | | Allometric coefficient for fine fuel shrub biomass (dry weight) |
| b_bsh | | | Allometric coefficient for fine fuel shrub biomass (dry weight) |
| a_btsh | | | Allometric coefficient for total fuel shrub biomass (dry weight) |
| b_btsh | | | Allometric coefficient for total fuel shrub biomass (dry weight) |
| cr | | [0-1] | Proportion of total height corresponding to the crown (i.e. Crown length divided by total height) |
| a_fbt | 0.041283131 | | Regression coefficient for tree foliar biomass |
| b_fbt | 1.415778672 | | Regression coefficient for tree foliar biomass |
| c_fbt | -0.016666626 | | Regression coefficient for tree foliar biomass |
| d_fbt | 0 | | Regression coefficient for tree foliar biomass |
| a_cr | 3.85373 | | Regression coefficient for crown ratio |
| b_1cr | -0.743 | | Regression coefficient for crown ratio |
| b_2cr | -0.02468 | | Regression coefficient for crown ratio |
| b_3cr | 0.000135686 | | Regression coefficient for crown ratio |
| c_1cr | 0 | | Regression coefficient for crown ratio |
| c_2cr | -0.38434 | | Regression coefficient for crown ratio |
| a_cw | 0.838617983 | | Regression coefficient for crown width |
| b_cw | 0.7347 | | Regression coefficient for crown width |
| LeafDuration | 0.667213248 | years | Duration of leaves in year |
| t0gdd | 98.4 | days | Date to start the accumulation of degree days |
| Sgdd | 201.9 | °C | Degree days for leaf budburst |
| Tbgdd | 0 | °C | Base temperature for the calculation of degree days to leaf budburst |
| Ssen | 5160 | °C | Degree days corresponding to senescence |
| Phsen | 12.5 | hours | Photoperiod corresponding to start counting senescence |

| | | | |
|---|---|---|---|
| Tbsen | 25 | °C | Base temperature for the calculation of degree days to senescence |
| xsen | 2 | {0,1,2} | Discrete values, to allow for any absent/proportional/more than proportional effects of temperature on senescence |
| ysen | 2 | {0,1,2} | Discrete values, to allow for any absent/proportional/more than proportional effects of photoperiod on senescence |
| SLA | 18.32 | $m^2kg^{-1}$ | Specific leaf area (mm2/mg = m2/kg) |
| LeafDensity | 0.462852877 | g $cm^{-3}$ | Density of leaf tissue (dry weight over volume) |
| WoodDensity | 0.679402099 | g $cm^{-3}$ | Wood tissue density (at 0% humidity!) |
| FineRootDensity | | g $cm^{-3}$ | Density of fine root tissue (dry weight over volume). |
| conduit2sapwood | 0.773166667 | [0,1] | Proportion of sapwood corresponding to conducive elements (vessels or tracheids) as opposed to parenchymatic tissue. |
| r635 | 2.302557615 | >=1 | Ratio of foliar (photosynthetic) + small branches (<6.35 mm) dry biomass to foliar (photosynthetic) dry biomass |
| pDead | | [0,1] | Proportion of total fine fuels that are dead |
| Al2As | 2076.12027 | $m^2 \cdot m^{-2}$ | Leaf area to sapwood area ratio |
| Ar2Al | | $m^2 \cdot m^{-2}$ | Root area to leaf area ratio |
| LeafWidth | 5.1177222 | cm | Leaf width |
| SRL | 522.1009957 | cm $g^{-1}$ | Specific root length |
| RLD | | cm $cm^{-3}$ | Fine root length density (density of root length per soil volume) |
| maxFMC | 113.5109237 | % | Maximum fuel moisture (in percent of dry weight) |
| minFMC | 86.19859579 | % | Minimum fuel moisture (in percent of dry weight) |
| LeafPI0 | -1.72 | Mpa | Osmotic potential at full turgor of leaves |
| LeafEPS | 11.89 | Mpa | Modulus of elasticity (capacity of the cell wall to resist changes in volume in response to changes in turgor) of leaves |
| LeafAF | 0.25 | % | Apoplastic fraction (proportion of water outside the living cells) in leaves |

| | | | |
|---|---|---|---|
| StemPI0 | | Mpa | Osmotic potential at full turgor of symplastic xylem tissue |
| StemEPS | | Mpa | Modulus of elasticity (capacity of the cell wall to resist changes in volume in response to changes in turgor) of symplastic xylem tissue |
| StemAF | | % | Apoplastic fraction (proportion of water outside the living cells) in stem xylem |
| SAV | | $m^2 m^{-3}$ | Surface-area-to-volume ratio of the small fuel (1h) fraction (leaves and branches < 6.35mm) |
| HeatContent | | $kJ\ kg^{-1}$ | High fuel heat content |
| LigninPercent | 12.56922096 | % | Percent of lignin+cutin over dry weight in leaves |
| gammaSWR | | unitless | Reflectance (albedo) coefficient for SWR (gammaPAR is 0.8*gammaSWR) |
| alphaSWR | | unitless | Absorbance coefficient for SWR (alphaPAR is alphaSWR*1.35) |
| kPAR | | unitless | Light extinction coeficient for PAR (extinction coefficient for SWR is kPAR/1.35) |
| g | | $mm\ LAI^{-1}$ | Canopy water storage capacity per LAI unit |
| Tmax_LAI | 0.130326956 | | Empirical coefficient relating LAI with the ratio of maximum transpiration over potential evapotranspiration. |
| Tmax_LAIsq | -0.005835535 | | Empirical coefficient relating squared LAI with the ratio of maximum transpiration over potential evapotranspiration. |
| Psi_Extract | -0.628424344 | Mpa | Water potential corresponding to 50% reduction of transpiration |
| Exp_Extract | 1.382918435 | | Parameter of the Weibull function regulating transpiration reduction |
| WUE | 7.924008761 | $g\ C \cdot mm\ H_2O^{-1}$ | Water use efficiency (gross photosynthesis over transpiration) |
| WUE_par | 0.321607784 | | Exponent regulating gross photosynthesis dependency on % PAR |
| WUE_co2 | 0.002396495 | | Exponent regulating gross photosynthesis dependency on atmospheric $CO_2$ concentration |
| WUE_vpd | -0.443200113 | | Exponent regulating gross photosynthesis dependency on atmospheric |

| | | | |
|---|---|---|---|
| | | | vapor pressure deficit |
| Gswmin | 0.004473333 | mol $H_2O \cdot s^{-1} \cdot m^{-2}$ | Minimum stomatal conductance to water vapour |
| Gswmax | 0.3 | mol $H_2O \cdot s^{-1} \cdot m^{-2}$ | Maximum stomatal conductance to water vapour |
| VCleaf_kmax | 8 | mmol $H_2O \cdot s^{-1} \cdot m^{-2} \cdot MPa^{-1}$ | Maximum leaf hydraulic conductance |
| VCleaf_c | 1.873130462 | | Parameter c of the leaf vulnerability curve |
| VCleaf_d | -1.737082162 | Mpa | Parameter d of the leaf vulnerability curve |
| Kmax_stemxylem | 0.9 | kg $H_2O \cdot s^{-1} \cdot m^{-1} \cdot MPa^{-1}$ | Maximum sapwood-specific hydraulic conductivity of stem xylem |
| VCstem_c | 7.314708 | | Parameter c of the stem xylem vulnerability curve |
| VCstem_d | -3.311856 | Mpa | Parameter d of the stem xylem vulnerability curve |
| Kmax_rootxylem | | kg $H_2O \cdot s^{-1} \cdot m^{-1} \cdot MPa^{-1}$ | Maximum sapwood-specific hydraulic conductivity of root xylem |
| VCroot_c | 1.873130462 | | Parameter c of the root xylem vulnerability curve |
| VCroot_d | -1.047264291 | Mpa | Parameter d of the root xylem vulnerability curve |
| Vmax298 | 94.5 | mmol $CO_2 \cdot s^{-1} \cdot m^{-2}$ | Maximum Rubisco carboxilation rate |
| Jmax298 | 159.9 | mmol electrons$\cdot s^{-1} \cdot m^{-2}$ | Maximum rate of electron transport at 298K |
| Nleaf | 23.45616703 | mg N g $dry^{-1}$ | Nitrogen mass per leaf dry mass |
| Nsapwood | 8.875 | mg N g $dry^{-1}$ | Nitrogen mass per sapwood dry mass |
| Nfineroot | 11.511125 | mg N g $dry^{-1}$ | Nitrogen mass per fine root dry mass |
| WoodC | 0.480582857 | g C g $dry^{-1}$ | Wood carbon content per dry mass |
| RERleaf | 0.021198634 | g gluc • g $dry^{-1}$ • $day^{-1}$ | Maintenance respiration rates for leaves. |
| RERsapwood | | g gluc • g $dry^{-1}$ • $day^{-1}$ | Maintenance respiration rates for living cells of sapwood. |

| | | | |
|---|---|---|---|
| RERfineroot | | g gluc • g dry$^{-1}$ • day$^{-1}$ | Maintenance respiration rates for fine roots. |
| CCleaf | 1.473 | g gluc • g dry$^{-1}$ | Leaf construction costs |
| CCsapwood | | g gluc • g dry$^{-1}$ | Sapwood construction costs |
| CCfineroot | | g gluc • g dry$^{-1}$ | Fine root construction costs |
| RGRleafmax | | m$^{2}$ cm$^{-2}$ day$^{-1}$ | Maximum leaf relative growth rate |
| RGRsapwoodmax | | cm$^{2}$ cm$^{-2}$ day$^{-1}$ | Maximum sapwood growth rate relative to sapwood area (for shrubs) |
| RGRcambiummax | 0.002672505 | cm$^{2}$ cm$^{-2}$ day$^{-1}$ | Maximum sapwood growth rate relative to cambium perimeter (for trees) |
| RGRfinerootmax | | g dry g dry$^{-1}$day$^{-1}$ | Maximum fineroot relative growth rate |
| SRsapwood | | day$^{-1}$ | Sapwood daily senescence rate |
| SRfineroot | | day$^{-1}$ | Fine root daily senescence rate |
| RSSG | 0.95 | [0-1] | Minimum relative starch for sapwood growth |
| MortalityBaselineRate | 0.0016 | year$^{-1}$ | Deterministic proportion or probability specifying the baseline reduction of cohort's density occurring in a year |
| SeedProductionHeight | | cm | Minimum height for seed production |
| ProbRecr | 0.01923847 | [0-1] | Probability of recruitment within the bioclimatic envelope |
| MinTempRecr | -4.66527874 | °C | Minimum average temperature of the coldest month for successful recruitment |
| MinMoistureRecr | 0.312279158 | unitless | Minimum value of the moisture index (annual precipitation over annual PET) for successful recruitment |
| MinFPARRecr | 0.193570197 | % | Minimum percentage of PAR at the ground level for successful recruitment |
| RecrTreeDBH | | cm | Recruitment tree dbh |
| RecrTreeHeight | 840.0000095 | cm | Recruitment tree height |

| | | | |
|---|---|---|---|
| RecrShrubHeight | | cm | Recruitment shrub height |
| RecrTreeDensity | 127.3239546 | Ind $ha^{-1}$ | Recruitment tree density |
| RecrShrubCover | | % | Recruitment shrub cover |
| RecrZ50 | | mm | Recruitment depth corresponding to 50% of fine roots |
| RecrZ95 | | mm | Recruitment depth corresponding to 95% of fine roots |

| 3D-CMCC-FEM parameters | | | |
|---|---|---|---|
| **Parameter Name** | **Value** | **Unit** | **Description** |
| LIGHT_TOL | 1 | unitless | 4 = very shade intolerant (cc = 90%), 3 = shade intolerant (cc = 100%), 2 = shade tolerant (cc = 110%), 1 = very shade tolerant (cc = 110%) |
| PHENOLOGY | 0.1 | unitless | 0.1 = deciduous broadleaf, 0.2 = deciduous needle leaf, 1.1 = broad leaf evergreen, 1.2 = needle leaf evergreen |
| ALPHA | 0.057 | mol C mol $PAR^{-1}$ | Canopy quantum efficiency |
| EPSILONgCMJ | 0.69 | g C $MJ^{-1}$ | Light Use Efficiency |
| GAMMA_LIGHT | 0 | unitless | Empirical parameter for Light modifiers |
| K | 0.71 | ratio | Extinction coefficient for absorption of PAR by canopy |
| ALBEDO | 0.13 | ratio | Albedo (varying from 0.13-0.17) |
| INT_COEFF | 0.3 | ratio | Precipitation interception coefficient |
| SLA_AVG0 | 20 | $m^2$ Kg $dry^{-1}$ | Average Specific Leaf Area (juvenile) sunlit/shaded leaves |
| SLA_AVG1 | 20 | $m^2$ Kg $dry^{-1}$ | Average Specific Leaf Area (mature) sunlit/shaded leaves |
| TSLA | 35 | year | Age at which SLA_AVG = (SLA_AVG1 + SLA_AVG0 )/2 |
| SLA_RATIO | 2.3 | ratio | Ratio of shaded to sunlit projected SLA |
| LAI_RATIO | 2 | ratio | All-sided to projected leaf area ratio |

| | | | |
|---|---|---|---|
| FRACBB0 | 0.125 | ratio | Branch and Bark fraction at age 0 |
| FRACBB1 | 0.125 | ratio | Branch and Bark fraction for mature stands |
| TBB | 20 | year | Age at which fracBB = (FRACBB0 + FRACBB1 )/ 2 |
| RHO0 | 0.64 | t dry $m^{-3}$ | Minimum Basic Density for young Trees |
| RHO1 | 0.64 | t dry $^{-3}$ | Maximum Basic Density for young Trees |
| TRHO | 100 | year | Age at which rho = (RHOMIN + RHOMAX )/2 |
| FORM_FACTOR | 0.433 | ratio | Define the ratio of the stem-volume and the cylinder volume defined by the tree height and tree BA |
| COEFFCOND | 0.08 | mbar | Define stomatal response to VPD |
| BLCOND | 0.01 | m $sec^{-1}$ | Canopy Boundary Layer conductance |
| MAXCOND | 0.006 | m $sec^{-1}$ | Maximum Stomatal Conductance |
| CUTCOND | 6.00E-05 | m $sec^{-1}$ | Cuticular conductance |
| MAXAGE | 400 | year | Determines rate of physiological declines of forest |
| RAGE | 0.95 | unitless | Relative Age to give fAGE = 0.5 |
| NAGE | 10 | unitless | Power of relative Age in function for Age |
| GROWTHTMIN | 0 | °C | Minimum temperature for growth 5 |
| GROWTHTMAX | 40 | °C | Maximum temperature for growth |
| GROWTHTOPT | 20 | °C | Optimum temperature for growth |
| GROWTHSTART | 60 | °C | Average temperature or (GDD) thermic sum for starting growth with Tbase=5°C |
| TAU | 4 | | Cold acclimation parameter (evergreen) |
| X0 | -2 | | Cold acclimation parameter (evergreen) |
| Smax | 9 | | Cold acclimation parameter (evergreen) |
| MINDAYLENGTH | 12 | ratio | Minimum day length |
| SWPOPEN | -0.34 | MPa | Leaf water potential: start of reduction |

| | | | |
|---|---|---|---|
| SWPCLOSE | -2.2 | MPa | Leaf water potential: complete reduction |
| OMEGA_CTEM | 0.8 | unitless | Allocation parameter control the sensitivity of allocation to changes in water and light availability |
| S0CTEM | 0.3 | ratio | Parameter controlling allocation to stem/Minimum ratio to Carbon to stem |
| R0CTEM | 0.3 | ratio | Parameter controlling allocation to root/Minimum ration to Carbon to roots |
| F0CTEM | 0.4 | ratio | Parameter controlling allocation to leaves |
| FRUIT_PERC | 0.05 | ratio | Fraction of NPP allocated for reproduction during the prescribed seasonal period |
| CONES_LIFE_SPAN | 0 | year | Life span for cones |
| FINE_ROOT_LEAF | 1.2 | ratio | Allocation new fine root C:new leaf |
| STEM_LEAF | 3.8 | ratio | Allocation new stem C:new leaf |
| COARSE_ROOT_STEM | 0.36 | ratio | Allocation new coarse root C:new stem |
| LIVE_TOTAL_WOOD | 0.13 | ratio | new live C:new total wood |
| N_RUBISCO | 0.1 | ratio | Fraction of leaf N in Rubisco |
| CN_LEAVES | 27 | kgC $kgN^{-1}$ | CN of leaves |
| CN_FALLING_LEAVES | 44 | kgC $kgN^{-1}$ | CN of leaf litter |
| CN_FINE_ROOTS | 72 | kgC $kgN^{-1}$ | CN of fine roots |
| CN_LIVEWOOD | 70 | kgC $kgN^{-1}$ | CN of live woods |
| CN_DEADWOOD | 550 | kgC $kgN^{-1}$ | CN of dead woods |
| LEAF_LITT_LAB_FRAC | 0.12 | ratio | Leaf litter labile fraction |
| LEAF_LITT_CEL_FRAC | 0.56 | ratio | Leaf litter cellulose fraction |
| LEAF_LITT_LIGN_FRAC | 0.32 | ratio | Leaf litter lignin fraction |
| FROOT_LITT_LAB_FRAC | 0.3 | ratio | Fine root litter labile fraction |
| FROOT_LITT_CEL_FRAC | 0.45 | ratio | Fine root litter cellulose fraction |

| | | | |
|---|---|---|---|
| FROOT_LITT_LIGN_FRAC | 0.25 | ratio | Fine root litter lignin fraction |
| DEADWOOD_CEL_FRAC | 0.75 | ratio | Dead wood litter cellulose fraction |
| DEADWOOD_LIGN_FRAC | 0.25 | ratio | Dead wood litter lignin fraction |
| BUD_BURST | 20 | days | Days of bud burst at the beginning of growing season |
| LEAF_FALL_FRAC_GROWING | 0.25 | ratio | Proportion of the growing season of leaf fall |
| LEAF_FINEROOT_TURNOVER | 1 | year | Average yearly fine root turnover rate |
| LIVEWOOD_TURNOVER | 0.02 | year | Annual yearly livewood turnover rate |
| SAPWOOD_TURNOVER | 0.02 | year | Annual yearly sapwood turnover rate |
| DBHDCMAX | 0.5 | ratio | Diameter/Canopy ratio at low Density |
| DBHDCMIN | 0.14 | ratio | Diameter/Canopy ratio at high Density |
| SAP_A | 0.391 | unitless | a parameter |
| SAP_B | 2.071 | unitless | b parameter |
| SAP_LEAF | 3310 | ratio | Leaf area/sapwood max ratio in pipe model |
| SAP_WRES | 0.11 | g TNC 100 g dry $^{-1}$ | Sapwood-Reserve biomass ratio |
| STEMCONST_P | 0.2837 | unitless | a parameter |
| STEMPOWER_P | 2.134 | unitless | b parameter |
| CRA | 35 | m | Chapman-Richards Maximum height H_MAX |
| CRB | 0.038 | unitless | Chapman_Richards b parameter |
| CRC | 1.104 | unitless | Chapman_Richards c parameter |
| HDMAX_A | 323.6 | unitless | A parameter for height to base diameter ratio MAX |
| HDMAX_B | -0.368 | unitless | B parameter for Height (m) to Base diameter (m) ratio MAX |
| HDMIN_A | 100.9 | unitless | A parameter for height to base diameter ratio MIN |
| HDMIN_B | -0.243 | unitless | B parameter for height to base diameter ratio MIN |

| | | | |
|---|---|---|---|
| CROWN_FORM_FACTOR | 0 | unitless | Crown form factor (0 = cylinder, 1 = cone, 2 = sphere) |
| CROWN_A | 0.3 | unitless | Crown relationship with tree height |
| CROWN_B | 1 | unitless | Crown exponential with tree height (Fixed to 1 as in Sortie-ND) |

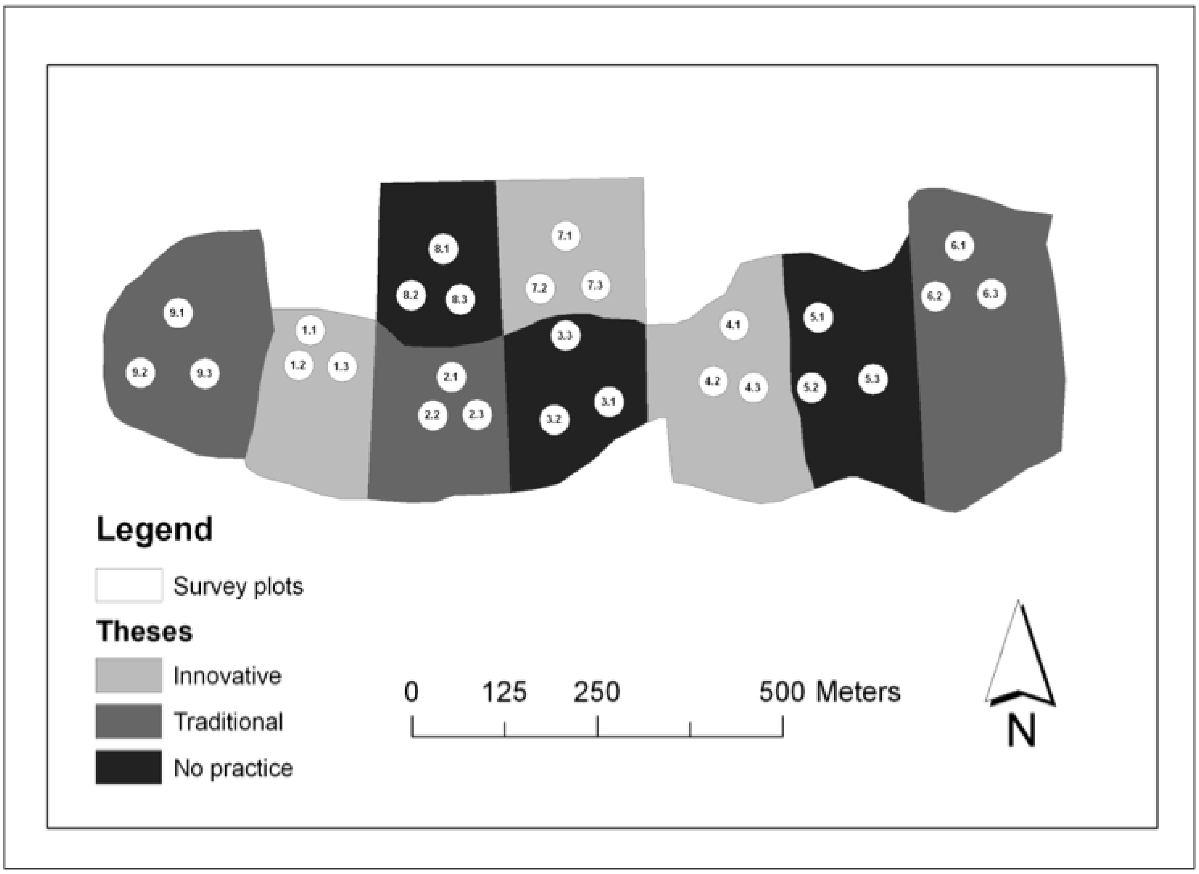


Fig S1. Layout design of Cansiglio experimental area (from https://www.manfor.eu/new/?page_id=379).

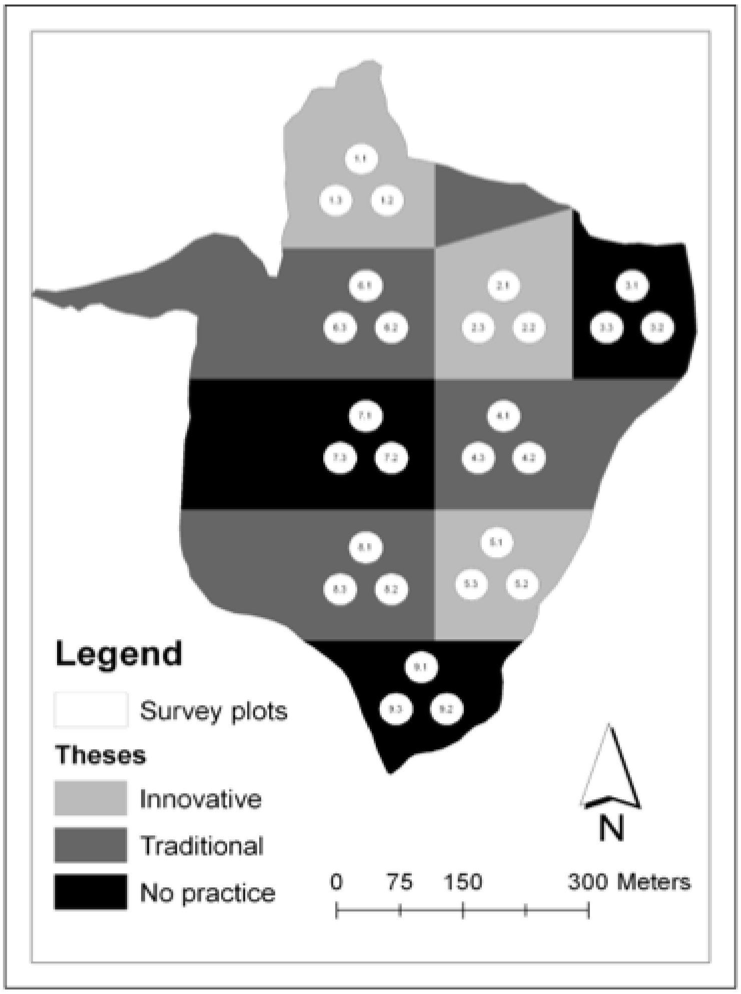


Fig S2. Layout design of Mongiana experimental area (from https://www.manfor.eu/new/?page_id=379).

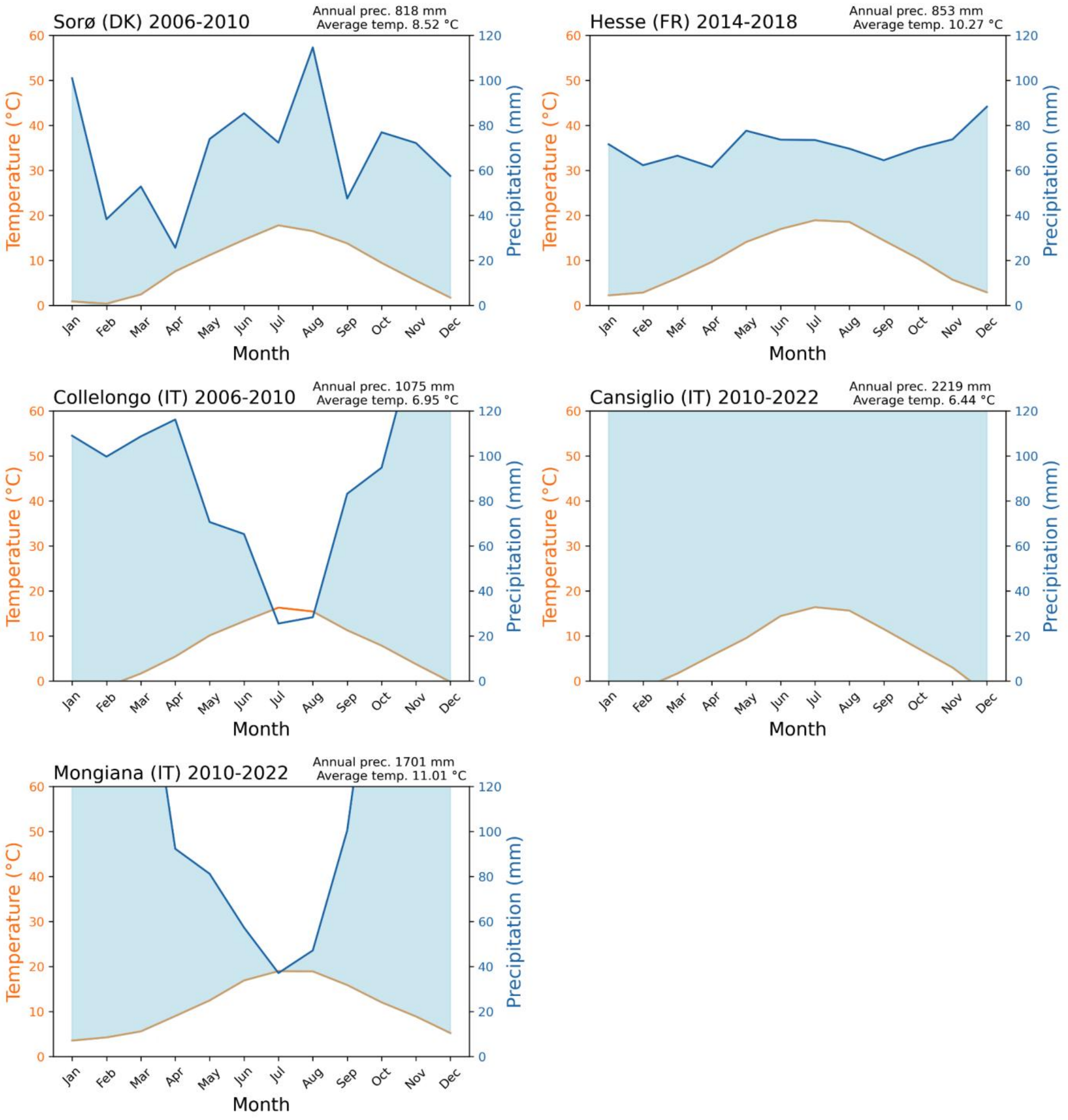


Fig S3. Bagnouls-Gaussen (Climatic diagram) of Sorø, Hesse, Collelongo, Cansiglio and Mongiana sites. Mean annual precipitation and mean annual temperature refers to the period indicated in the title.

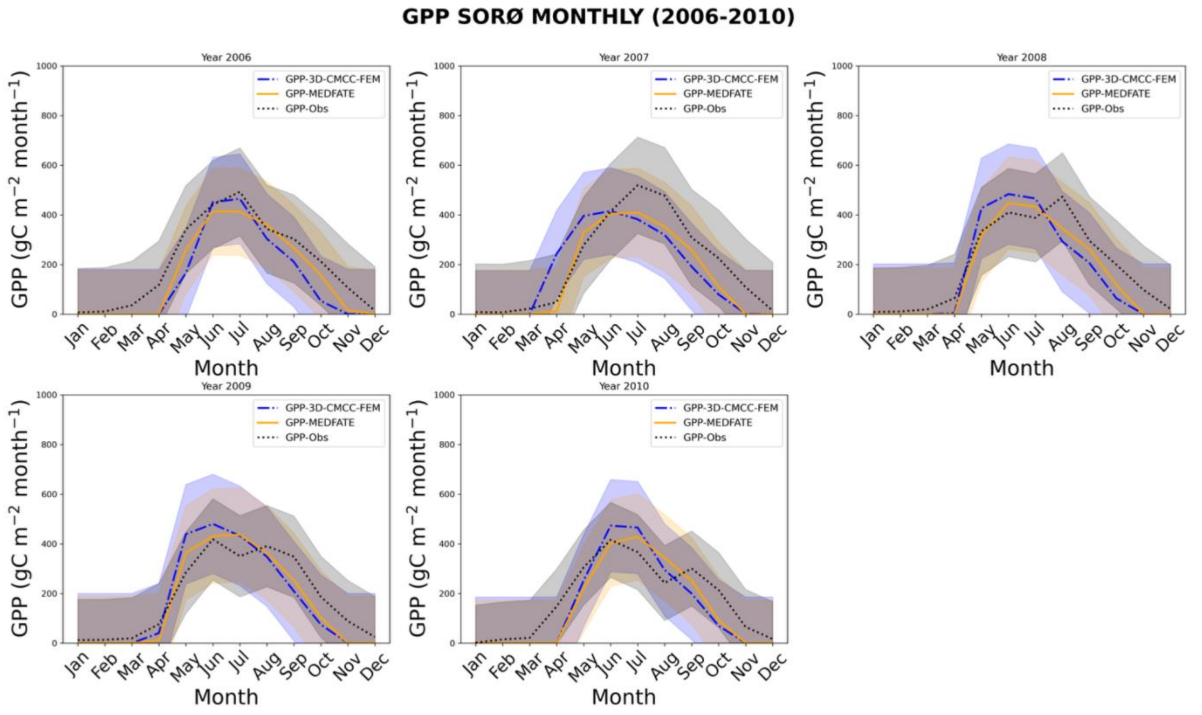


Fig S4. Monthly mean seasonal cycle of GPP (gC $m^{-2}$ $day^{-1}$) estimated from the direct micrometeorological eddy covariance measurements (GPP-Obs) and models' simulation (GPP-3D-CMCC-FEM and, GPP-MEDFATE) during the evaluation period at the DK-Sor at the Beech forest in 2006-2010.

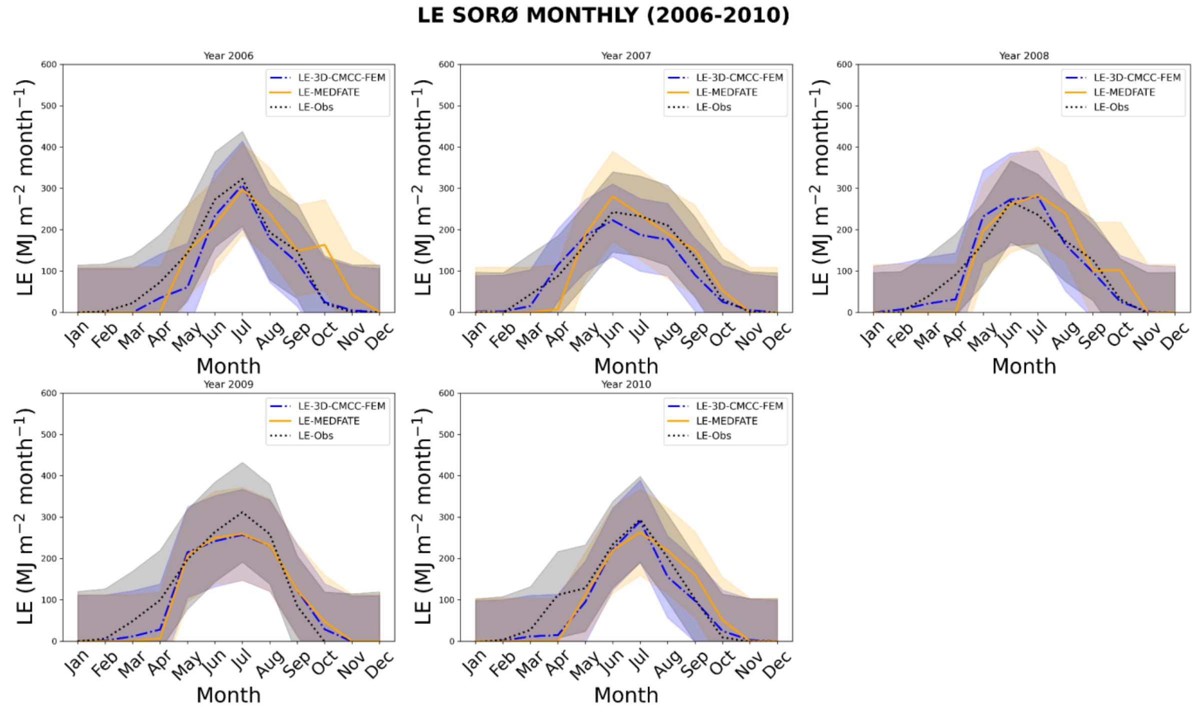


Fig S5. Monthly mean seasonal cycle of LE (MJ $m^{-2}$ $day^{-1}$) estimated from the direct micrometeorological eddy covariance measurements (LE-Obs) and models' simulation (LE-3D-CMCC-FEM and LE-MEDFATE) during the evaluation period at the DK-Sor at the Beech forest in 2006-2010.

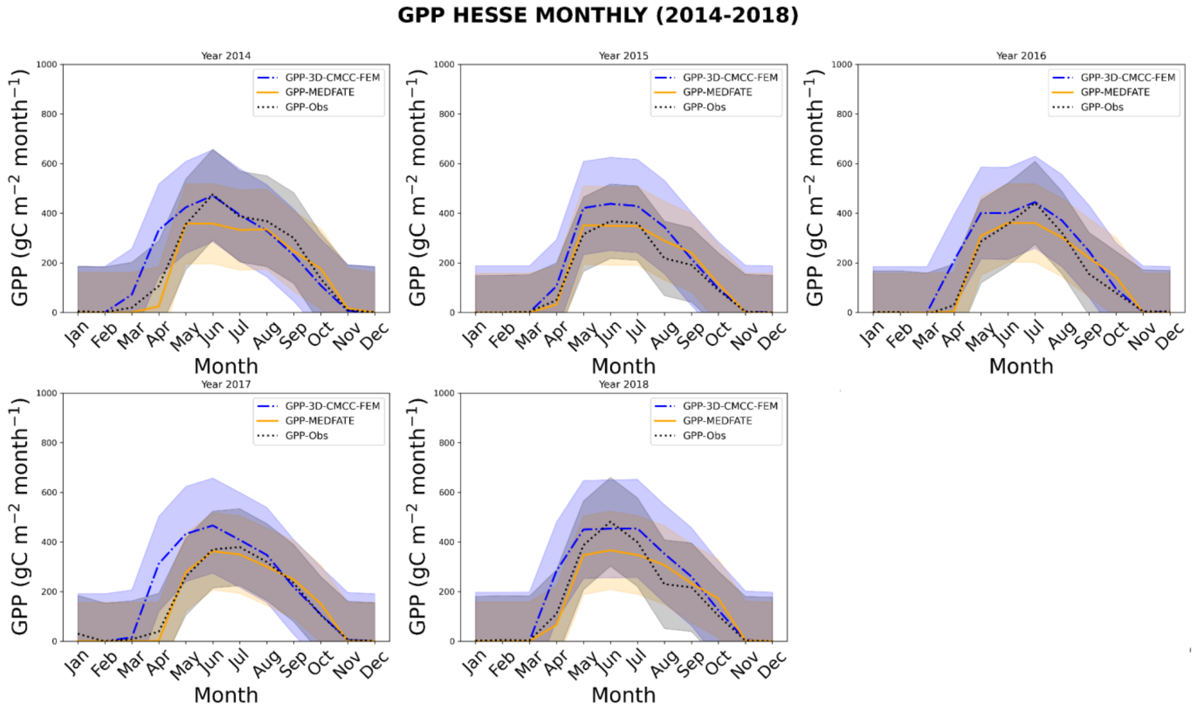


Fig S6. Monthly mean seasonal cycle of GPP (gC $m^{-2}$ $day^{-1}$) estimated from the direct micrometeorological eddy covariance measurements (GPP-Obs) and models' simulation (GPP-3D-CMCC-FEM and, GPP-MEDFATE) during the evaluation period at the FR-Hes at the Beech forest in 2014-2018.

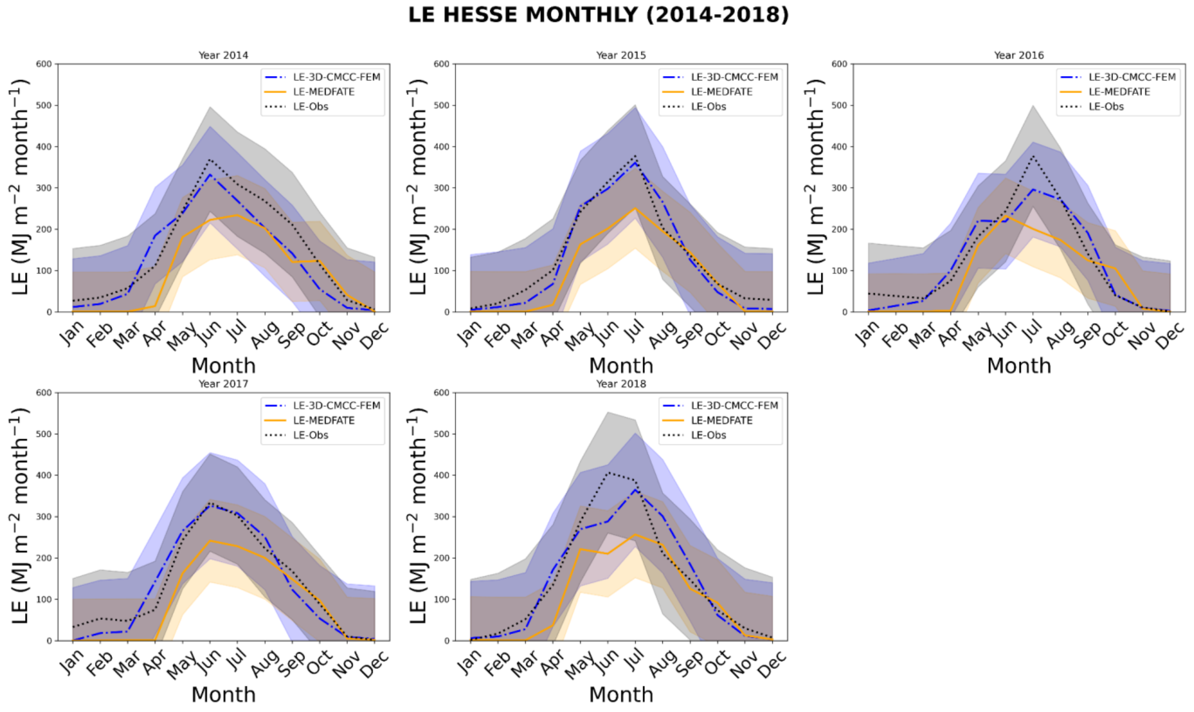


Fig S7. Monthly mean seasonal cycle of LE (MJ $m^{-2}$ $day^{-1}$) estimated from the direct micrometeorological eddy covariance measurements (LE-Obs) and models' simulation (LE-3D-

CMCC-FEM and LE-MEDFATE) during the evaluation period at the FR-Hes at the Beech forest in 2014-2018.

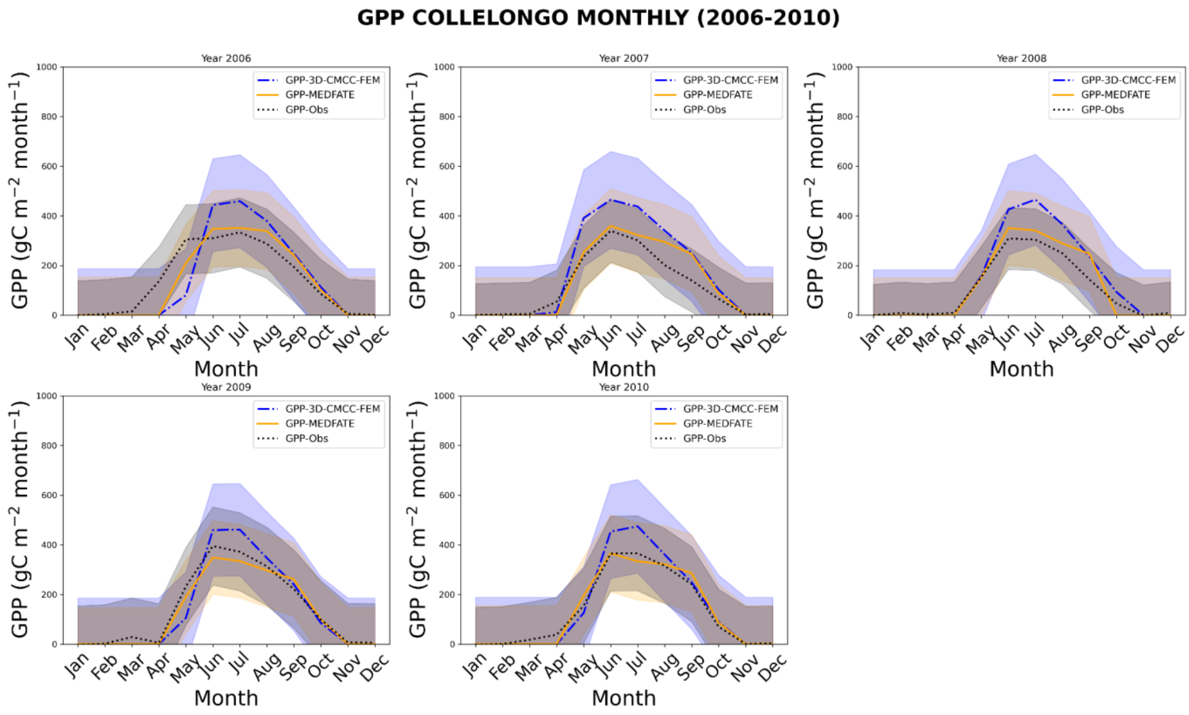


Fig S8. Monthly mean seasonal cycle of GPP (gC $m^{-2}$ $day^{-1}$) estimated from the direct micrometeorological eddy covariance measurements (GPP-Obs) and models' simulation (GPP-3D-CMCC-FEM and, GPP-MEDFATE) during the evaluation period at the IT-Col at the Beech forest in 2006-2010.

**LE COLLELONGO MONTHLY (2006-2010)**

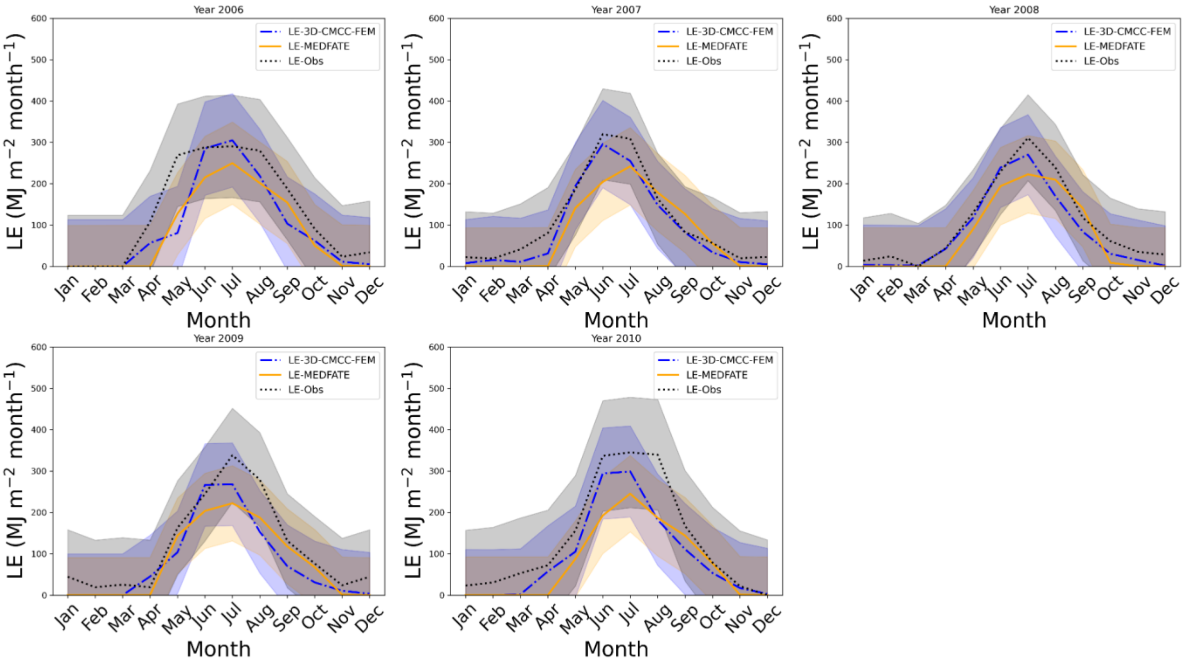


Fig S9. Monthly mean seasonal cycle of LE (MJ $m^{-2}$ $day^{-1}$) estimated from the direct micrometeorological eddy covariance measurements (LE-Obs) and models' simulation (LE-3D-CMCC-FEM and LE-MEDFATE) during the evaluation period at the IT-Col at the Beech forest in 2006-2010.

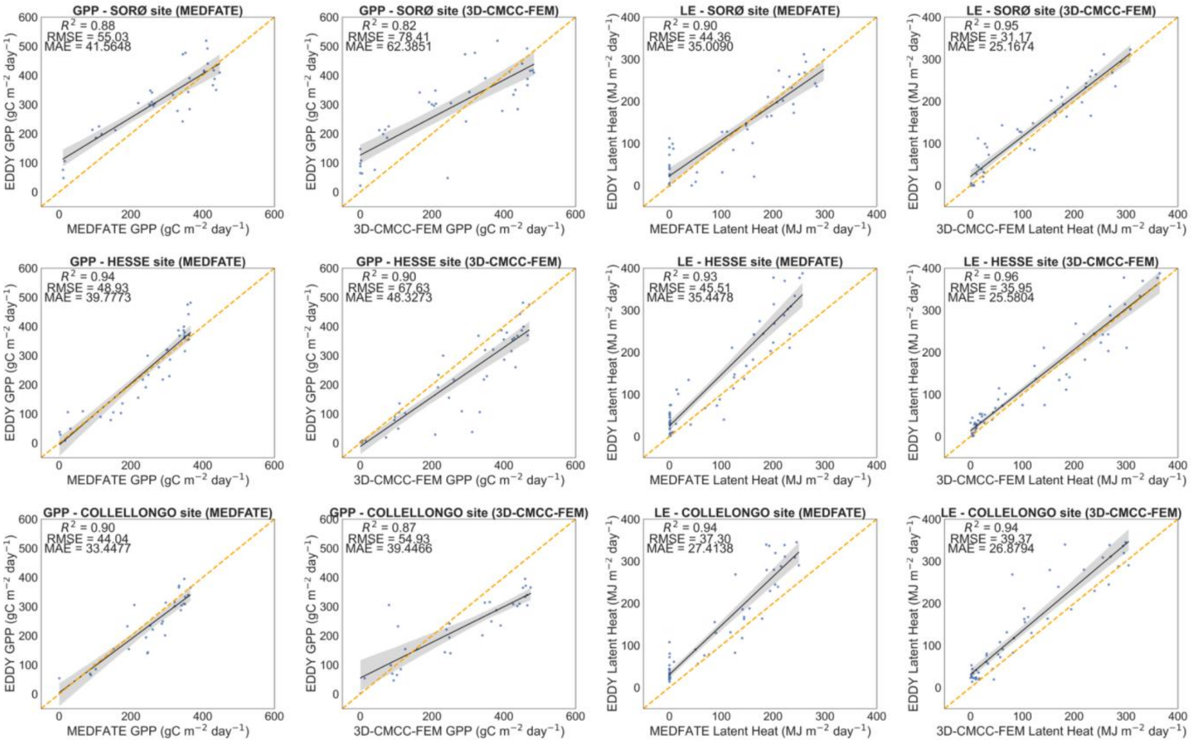


Fig S10. Scatter plots and linear regressions of GPP (gC $m^{-2}$ $day^{-1}$) and LE (MJ $m^{-2}$ $day^{-1}$) of the models versus the direct micrometeorological eddy covariance measurements (Obs) at the Sorø (DK-Sor; 2006-2010 period), Collelongo (IT-Col; 2006-2010 period) and Hesse (FR-Hes; 2014-2018 period).

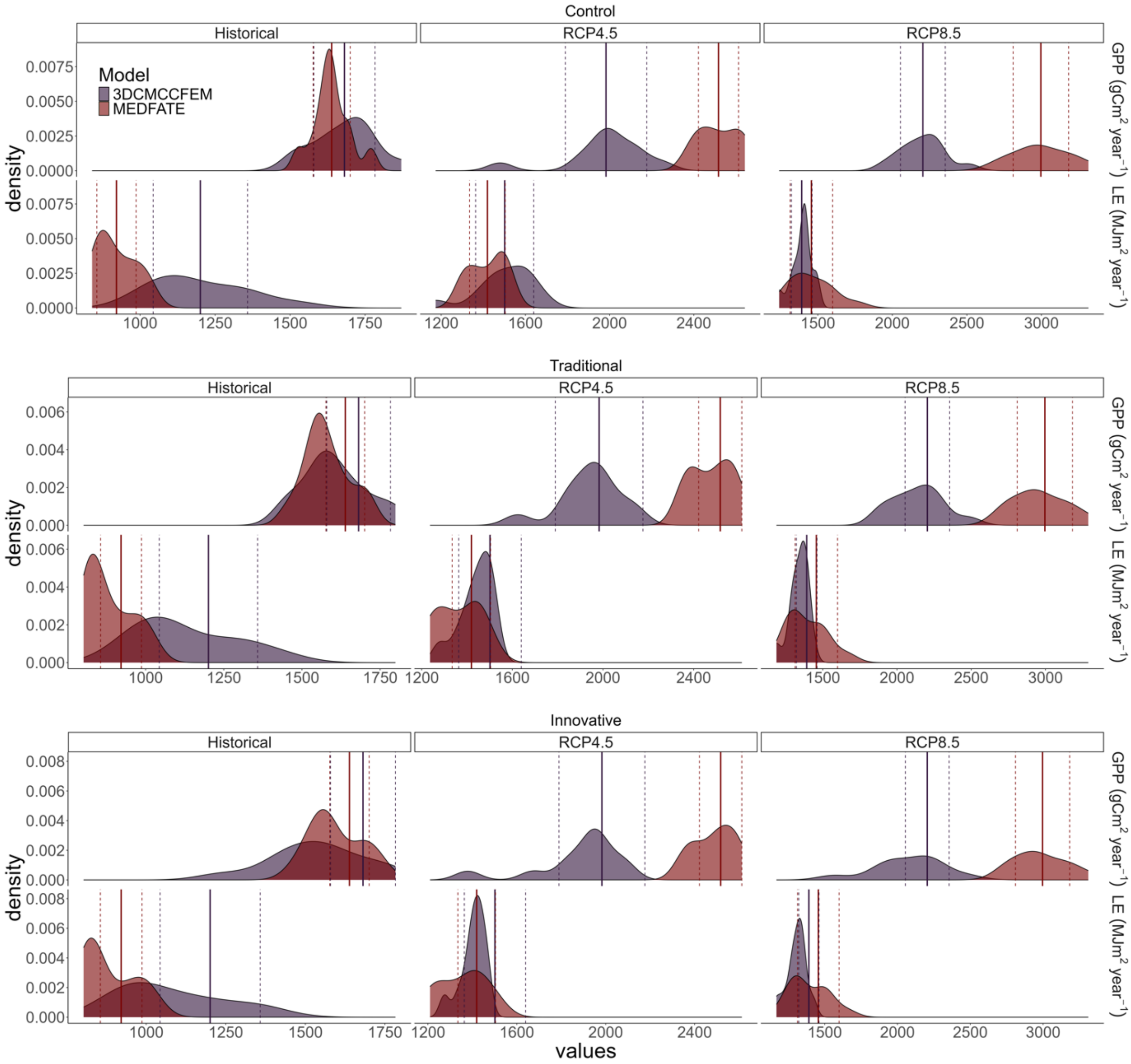


Fig S11. Comparative analysis between models output at the Cansiglio site. The figure is divided into three horizontal panels, each representing different management strategies for the plot: 'Control' (top panel), 'Traditional' (middle panel), and 'Innovative' (bottom panel). Within each panel, annual GPP (gC $m^{-2}$ $year^{-1}$) and annual LE (MJ $m^{-2}$ $yr^{-1}$) are displayed, as modeled by MEDFATE and 3D-CMCC-FEM, respectively. The columns represent different climate scenarios: historical data ('Hist', 2010–2022) on the left, projections under the RCP4.5 scenario (2059–2070) in the middle, and projections under the RCP8.5 scenario (2059–2070) on the right. The solid line represents the mean, while the dotted line indicates the standard deviation.

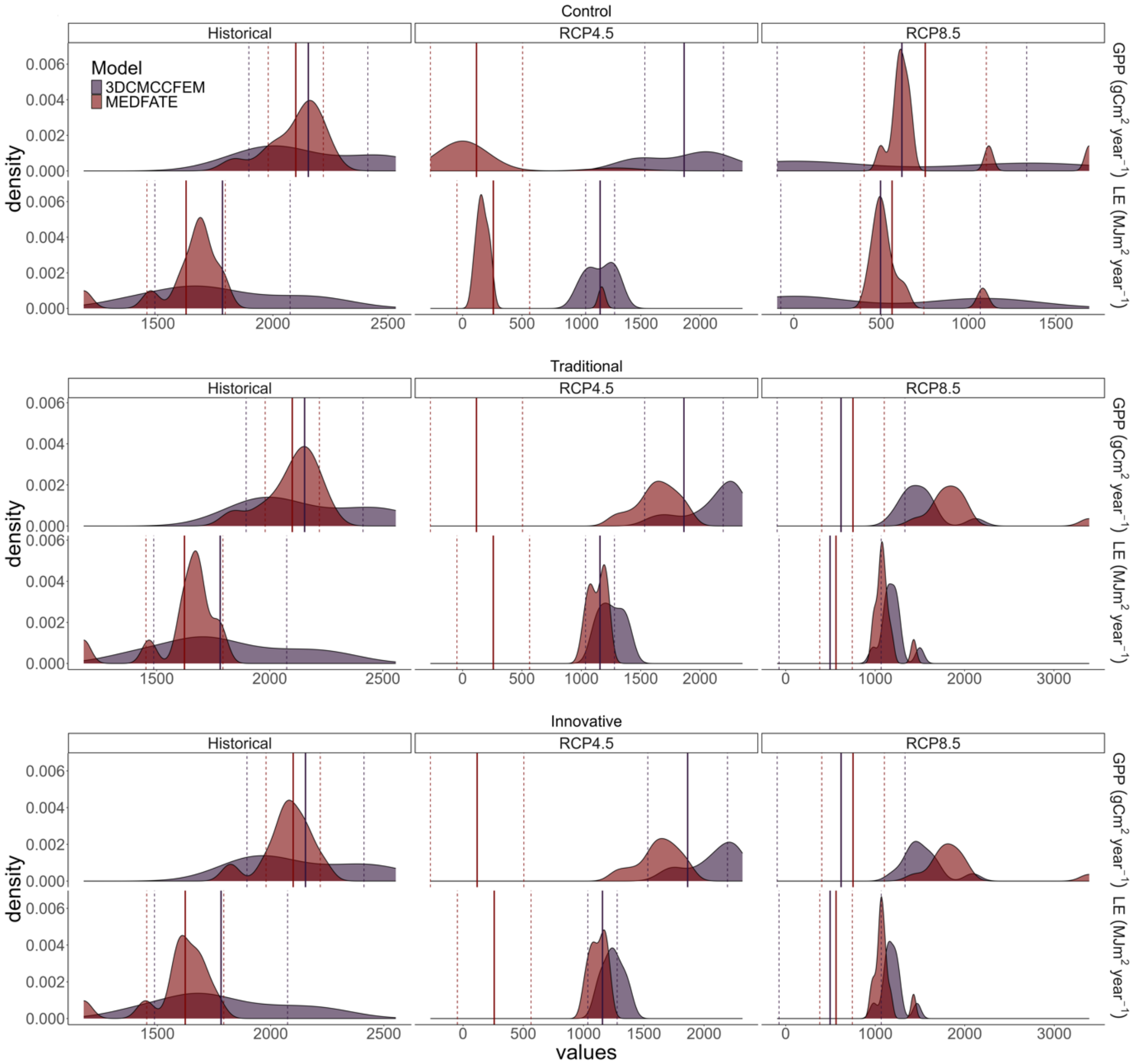


Fig S12. Comparative analysis between models output at the Mongiana site. The figure is divided into three horizontal panels, each representing different management strategies for the plot: 'Control' (top panel), 'Traditional' (middle panel), and 'Innovative' (bottom panel). Within each panel, annual GPP (gC $m^{-2}$ $year^{-1}$) and annual LE (MJ $m^{-2}$ $yr^{-1}$) are displayed, as modeled by MEDFATE and 3D-CMCC-FEM, respectively. The columns represent different climate scenarios: historical data ('Hist', 2010–2022) on the left, projections under the RCP4.5 scenario (2059–2070) in the middle, and projections under the RCP8.5 scenario (2059–2070) on the right. The solid line represents the mean, while the dotted line indicates the standard deviation.
